# Cholinergic interneuron control of GABAergic circuits targeting spiny projection neurons is disrupted in parkinsonian models

**DOI:** 10.64898/2026.06.17.732978

**Authors:** M. Belal, T. Perez-Rosello, E. B. Guven, S. Kocaturk, Z. Xie, E. Ilijic, T. Tkatch, J. Li, W. Dauer, M. Assous, J.M. Tepper, V. R. J. Clarke, D. J. Surmeier

**Affiliations:** Department of Neuroscience, Feinberg School of Medicine, Northwestern University, Chicago, IL 60611, USA; Aligning Science Across Parkinson’s (ASAP) Collaborative Research Network, Chevy Chase, MD 20815, USA; Molecular and Behavioral Neuroscience, Rutgers University, Newark, NJ, USA; School of Biosciences, Cardiff University, Cardiff, UK; Department of Internal Medicine, University of Michigan Medical School. Ann Arbor, MI, USA; Peter O’Donnell Jr. Brain Institute, Departments of Neurology and Neuroscience, University of Texas Southwestern Medical Center, Dallas, TX, USA

**Keywords:** interneuron, striatum, nicotinic, dendrite, dopamine, synapse, computational

## Abstract

Parkinson’s disease (PD) is known to alter the intrinsic properties of striatal cholinergic interneurons (ChIs). However, how PD shapes ChI control of intrastriatal GABAergic circuits regulating principal spiny projection neurons (SPNs) is unknown. To fill this gap, striatal circuits in healthy and parkinsonian mice were interrogated. In *ex vivo* brain slices from healthy mice, optogenetic stimulation of ChIs evoked GABA_A_ receptor currents in both indirect and direct pathway SPNs that were attributable to nicotinic acetylcholine receptor (nAChR)-mediated activation of GABAergic interneurons (GIs). Simulations suggested that this circuit exerts a state-dependent control of SPN dendritic integration that was modulated by concomitant muscarinic receptor signaling. Surprisingly, in mouse models of prodromal and parkinsonian states, the ability of ChIs to engage this intrastriatal circuitry was disrupted because interneurons down-regulated nicotinic AChRs. Taken together, these studies suggest that impaired ChI control of GABAergic interneurons contributes to behavioral deficits in both prodromal and clinical PD states.

## Introduction

The striatum is a key node in the brain circuitry controlling goal-directed action and habit ^1^. In the dorsal striatum, SPNs can be divided into two broad classes, indirect pathway SPNs (iSPNs) and direct pathway SPNs (dSPNs), on the basis of their intrinsic properties, functional connectivity, and role in movement control ^2,3^. The activity of iSPNs and dSPNs is governed by a robust extra-striatal innervation arising from the cerebral cortex, thalamus and other parts of the basal ganglia, including dopaminergic neurons in the substantia nigra pars compacta (SNc). In addition, SPNs are innervated by a diverse array of striatal interneurons, including giant ChIs and a diverse collection of GABAergic interneurons ^2,4^.

Although a small fraction of all striatal neurons, ChIs figure prominently in models of striatal function, being linked to set-shifting, reversal learning, and movement sequences ^5–11^. ChI control of the striatal circuitry relies upon both fast and slow mechanisms. Acting through G-protein coupled muscarinic acetylcholine receptors (mAChRs), ChIs differentially modulate the somatodendritic excitability of iSPNs and dSPNs and shape the information reaching them from corticostriatal pyramidal neurons ^12–14^. In addition to this slow, second messenger dependent modulation, ChIs rapidly excite tyrosine hydroxylase-expressing interneurons (THIs) and neurogliaform interneurons (NGFIs) through nicotinic acetylcholine receptors (nAChRs) ^15^; these two types of GABAergic interneuron target the dendrites of SPNs, in contrast to fast-spiking GABAergic interneurons, which preferentially target SPN perisomatic regions ^4,15^. Although they target similar subcellular sites, the postsynaptic currents (PSCs) evoked by THIs are transient and typical of synaptic GABA_A_ receptors (GABA_A_R), whereas those evoked by NGFIs are slower and have been attributed to extrasynaptic GABA_A_Rs, although definitive evidence for this assertion is lacking ^4,15^. It also is unclear whether the strength of this ChI-driven GABAergic circuit is similar in iSPNs and dSPNs. Given that the mAChR modulation of iSPN increases, whereas that of dSPN largely decreases excitability ^12^, one might expect ChI-driven GABAergic inhibition of dSPNs to be stronger. That said, GABA_A_R-mediated effects on SPNs cannot be viewed as simply inhibitory, as opening of GABA_A_Rs depolarizes ‘resting’ SPNs and can promote dendritic spikes ^16,17^, making a priori predictions about coupling strength less than straightforward.

In PD, the striatal circuitry is profoundly disrupted by the degeneration of SNc dopaminergic axons ^18,19^. In addition to interrupting the dopaminergic modulation of dSPNs and iSPNs, the loss of striatal dopamine (DA) in models of late-stage PD results in a hyper-cholinergic state ^20^. This pathological state is attributable to dis-inhibition of evoked acetylcholine (ACh) release, as well as alterations in the intrinsic properties of ChIs ^20–22^. But how changes in cholinergic signaling contribute to the defining motor deficits of PD is unclear. Recent work with a progressive mouse model of PD (MCI-Park) has shown that striatal DA depletion alone is not sufficient to trigger the broader network dysfunction necessary for the emergence of the motor disability characteristic of PD ^23^. In part, this departure from the classical model of network dysfunction underlying PD ^24^ could reflect the induction of homeostatic plasticity in the ChI-driven intrastriatal circuitry regulating iSPN and dSPN activity.

The experiments described here employed a combination of genetic, optical, electrophysiological and computational approaches in healthy, MCI-Park and 6-hydroxydopamine (6-OHDA) lesioned mice to gain a better understanding of ChI-controlled intrastriatal circuits. These studies revealed that in healthy mice, ChI-driven activity in THIs and NGFIs produced similar PSCs in iSPNs and dSPNs. Simulations revealed that in ‘resting’ or down-state iSPNs and dSPNs, this GABAergic signaling, particularly that through NGFIs, created a temporal window in which the ability of distal dendritic glutamatergic input to evoke local spikes was enhanced. Interestingly, in iSPN simulations, mAChR-mediated modulation of K+ channels worked in concert with the ChI-driven GABAergic circuit to promote down-state dendritic spikes, while minimizing the GABAergic suppression of somatic spike generation in response to suprathreshold glutamatergic inputs. In the MCI-Park model of prodromal PD, evoked ACh release was elevated, as in models of late-stage PD (6-OHDA). However, the ability of ChIs to drive GABAergic PSC in iSPNs and dSPNs was dramatically down-regulated in MCI-Park as well as in 6-OHDA lesioned mice. This uncoupling was attributable in part to a down-regulation in the expression of nAChRs in THIs and NGFIs, not to a disruption in the coupling of these GABAergic interneurons to SPNs. Although homeostatic, this uncoupling should impair the ability of ChIs to modulate transitions in the engagement of SPN ensembles, contributing to the behavioral deficits observed in both prodromal and parkinsonian stages of PD.

## Results

### Optogenetic stimulation of ChIs evokes a complex synaptic response in iSPNs and dSPNs

To investigate the coupling between ChIs and iSPNs and dSPNs, mice expressing Cre recombinase under the choline acetyltransferase (ChAT) promoter were crossed either with mice expressing tdTomato under the control of the *Drd1a* promoter (ChAT-Cre:: *Drd1a^tdTomato^*) or the *Drd2* promoter (ChAT-Cre::*Drd2^e^*^GFP^). To optogenetically manipulate the activity of ChIs, an adeno-associated virus (AAV) carrying a Cre recombinase-dependent Chronos expression construct was stereotaxically injected into the DLS of young adult mice (Fig. 1a). After allowing time for Chronos expression in ChIs (Fig. 1b), mice were sacrificed and brain slices prepared as described previously ^16^. In whole cell voltage clamp recordings from SPNs, optogenetic activation of ChIs evoked a multi-phasic synaptic response with fast and slow components (Fig. 1c, d). Bath application of the GABA_A_R antagonist gabazine (10 µM) profoundly reduced ChI-evoked PSCs (Fig. 1d). The amplitude and the area of the ChI-evoked PSCs were significantly reduced by gabazine in both iSPNs and dSPNs (Fig. 1e, f).

**Figure 1.**
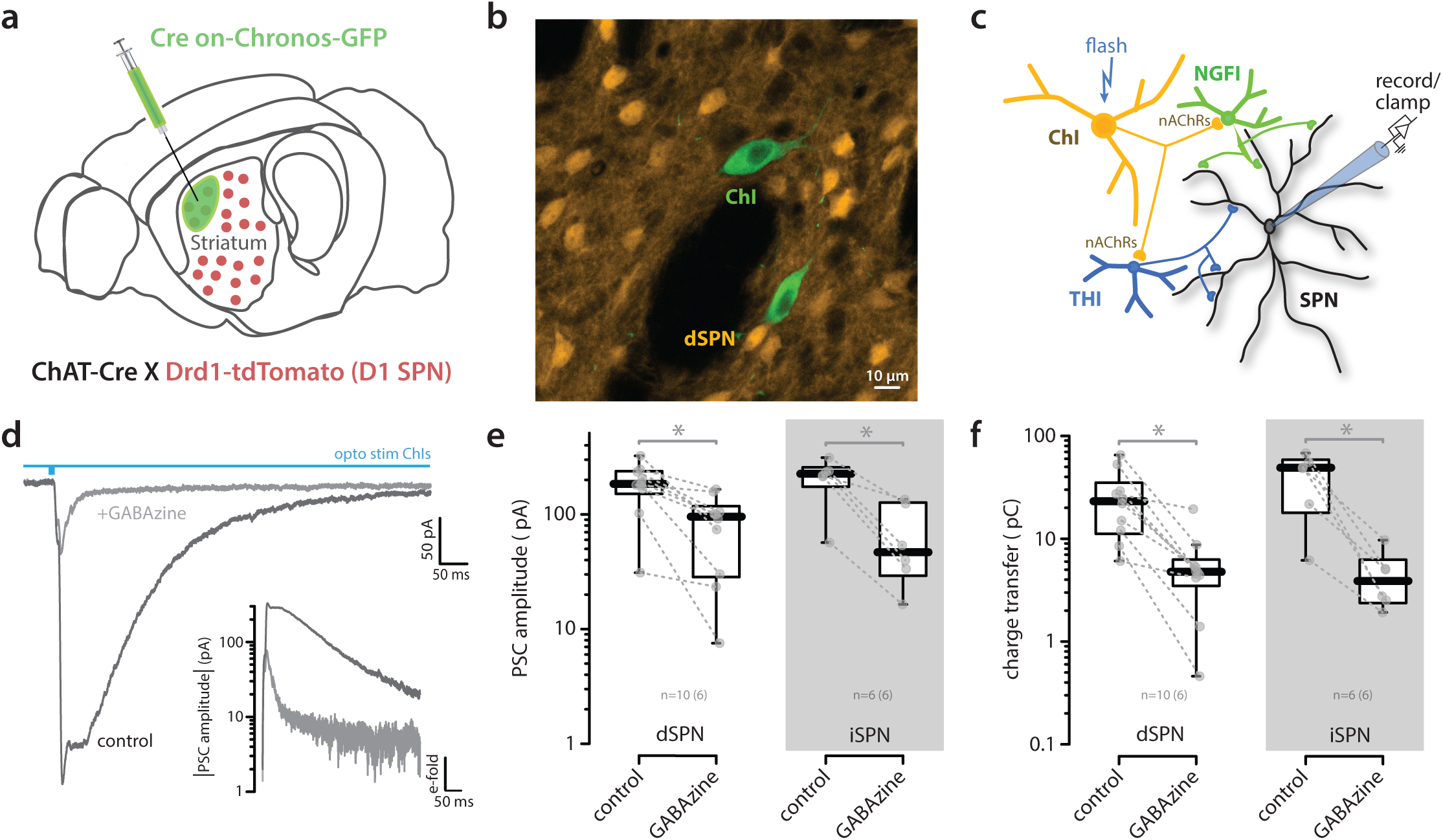
Bath application of gabazine reduced ChI-evoked PSCs in SPNs. (a) Schematic of viral injection of Chronos into the DLS of ChAT-Cre X Drd1-tdTomato mouse. (b) Confocal image of a coronal striatal section showing selective expression of Chronos in cholinergic interneurons (green) and tdTomato in dSPNs (orange). (c) Circuit diagram depicting the cholinergic control of GABAergic microcircuit targeting SPNs. (d) Representative PSCs recorded in a dSPN in response to activation of ChIs (whole-field LED, 5 ms) under control conditions and after bath application of gabazine (10 µM). (e) Semi-log box plots of PSC amplitudes in dSPNs (n = 10, 6 animals) and iSPNs (n = 6, 6 animals) showing reduced responses by gabazine (Wilcoxon signed-rank test: dSPN, V = 0, *p* = 0.00391; iSPN, V = 0, *p* = 0.03125). (f) Semi-log box plots of charge transfer for the same cells showing reduced charge transfer by gabazine (Wilcoxon signed-rank test: dSPN, V = 55, *p* = 0.00391; iSPN, V = 21, *p* = 0.03125). On this and subsequent figures, all relevant scale bars are illustrated and labeled in the corresponding panel.

As previously described ^15,25^, the gabazine-sensitive currents were biphasic (Fig. 1d). The fast component of the response has been attributed to nAChR-mediated activation of THIs and the slow component to nAChR-mediated activation of NGFIs ^15,25^ (Fig. 2a). To deconstruct the PSCs into components that could be nominally attributed to THIs and NGFIs, they were fitted with the sum of two functions, each modeled as the product of an exponential rise and an exponential decay (see Methods). Interestingly, the PSCs evoked in iSPNs and dSPNs were very similar (Fig. 2b-d). To assess the relative contribution of these components to the aggregate PSCs, the amplitudes of the fast and slow components in individual iSPNs and dSPNs were plotted (Fig. 2e). The absolute amplitude of the slow PSC typically was larger than that of the fast, nominally THI-mediated PSC in both iSPNs and dSPNs. Furthermore, the PSC decay time constants for the fast and slow components were similar in iSPNs and dSPNs (Fig. 2f).

**Figure 2.**
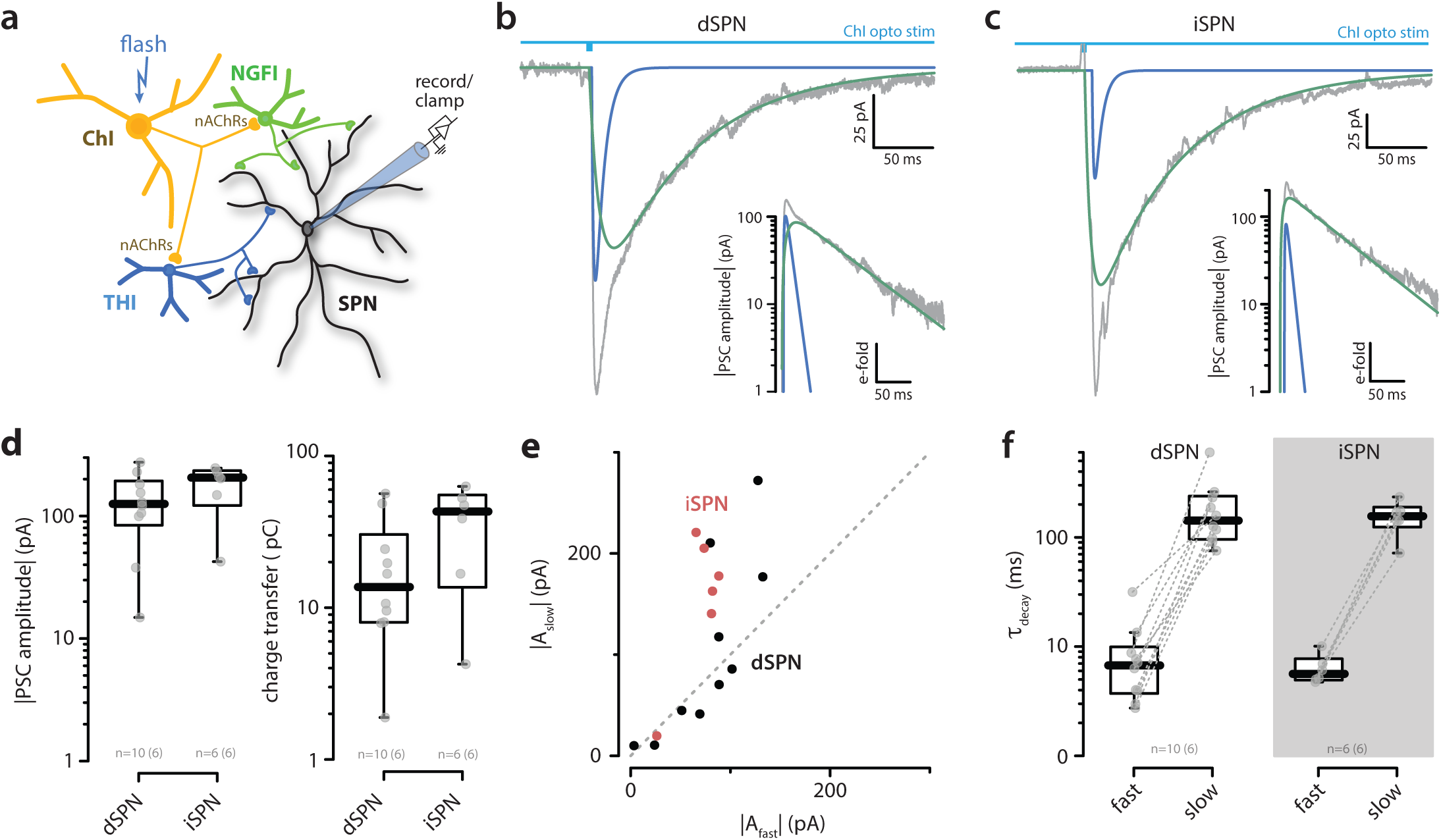
ChI-evoked PSCs were similar in iSPNs and dSPNs. (a) Circuit diagram depicting the cholinergic–GABAergic microcircuit onto SPNs. (b) Representative gabazine-sensitive PSC recorded from a dSPN and (c) iSPN in response to activation of ChIs (whole-field LED, 5 ms), composed of fast- and slow-decaying components. Inset, semi-logarithmic scale. (d) Semi-log box plots of PSC amplitudes and total charge transfer in dSPNs (n = 10, 6 animals) and iSPNs (n = 6, 6 animals) showing no difference between cell types (Wilcoxon rank-sum test: amplitude, W = 42, *p* = 0.21978; charge transfer, W = 18, *p* = 0.21978). (e) Scatter plot of slow-decaying versus fast-decaying amplitudes in dSPNs and iSPNs showing consistently larger slow-decaying responses. (f) Semi-log box plots of fitted decay time constants for fast- and slow-decaying components in dSPNs (n = 10, 6 animals) and iSPNs (n = 6, 6 animals) showing similar kinetics (Wilcoxon rank-sum test, dSPN vs iSPN: fast, W = 31, *p* = 1; slow, W = 30, *p* = 1).

### The slow component of ChI-evoked GABA_A_R currents included those mediated by extrasynaptic receptors

Previously, the slow kinetics of the NGFI-mediated PSC in SPNs has been attributed to the engagement of extrasynaptic GABA_A_Rs ^15^. But, this hypothesis has not been tested. Typically, extrasynaptic GABA_A_Rs differ in subunit composition from those found at synaptic sites: synaptic GABA_A_Rs contain a γ subunit along with α and β subunits, whereas extrasynaptic receptors contain δ subunits in combination with α4 or α6 subunits ^26^. To test the hypothesis that the slow component of the ChI-evoked PSC in SPNs was attributable to extrasynaptic GABA_A_Rs, the CRISPR/Cas9 gene-editing system was used to disrupt δ subunit expression in SPNs. Specifically, two AAV vectors, one carrying a Cre-independent Cas9 expression construct and another carrying two δ subunit guide RNAs (δ-gRNA) expression constructs, were injected along with an AAV carrying the Cre-dependent Chronos construct into the dorsolateral striatum (DLS) of a ChAT-Cre mouse (Fig. 3a). Four weeks later, mice were sacrificed and brain slices prepared for study. In these mice, the total charge transfer by ChI-evoked PSCs was reduced (Fig. 3b-d). With deconstruction of the two current components, it became clear that the amplitude of the slow but not the fast component of the PSC was reduced by δ subunit knockdown (Fig. 3e). However, the kinetics of the fast and slow components were not altered (Fig. 3f).

**Figure 3.**
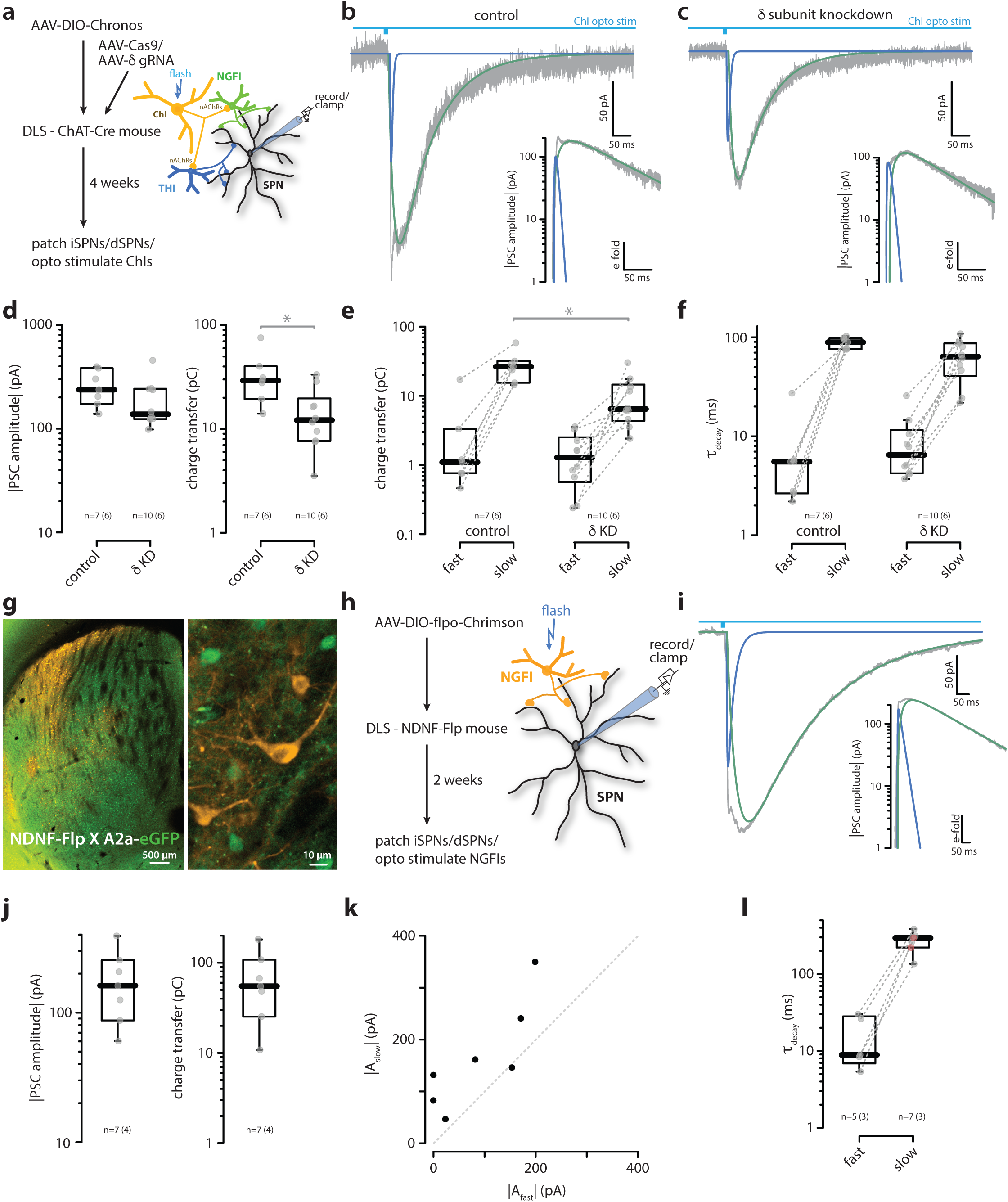
The slow component of ChI-evoked GABAergic currents was mediated in part by GABA_A_δ-subunits. (a) Experimental timeline and schematic of the ChI–SPN microcircuit. Representative PSC recorded from an SPN in response to activation of ChIs (whole-field LED, 5 ms) under control conditions (b) and following δ-subunit knockdown (c) showing fast- and slow-decaying components. Inset, semi-log scale. (d) Semi-log box plots of PSC amplitude and charge transfer in control (n = 7, 6 animals) and δ-subunit knockdown (n = 10, 6 animals) showing reduced total charge transfer with no change in maximum amplitude (Wilcoxon rank-sum test: amplitude, W = 19, *p* = 0.13307; charge transfer, W = 60, *p* = 0.01357). (e) Semi-log box plots of charge transfer for fast- and slow-decaying components in control and δ-subunit knockdown showing selective reduction of the slow component (Wilcoxon rank-sum test, control vs knockdown: fast, W = 23, *p* = 0.26985; slow, W = 62, *p* = 0.01357). (f) Semi-log box plots of decay time constants in control and δ-subunit knockdown showing no change in decay kinetics (Wilcoxon rank-sum test, control vs knockdown: fast, W = 18, *p* = 0.21761; slow, W = 40, *p* = 0.66907). (g-l) **The slow component of NGFI-evoked GABAergic currents in SPNs is kinetically similar to the Chi-evoked current**. (g) Confocal image of a striatal section showing tdTomato expression in NDNF-expressing NGF interneurons (orange) and A2a-eGFP expression in iSPNs (green). (h) Experimental timeline and schematic of the NGF–SPN microcircuit. (i) Representative PSC recorded from an SPN after activating NGF interneurons (whole-field LED, 5 ms). Inset, semi-logarithmic scale. (j) Semi-log box plots of PSC amplitude and charge transfer in SPNs after NGF interneuron activation. (k) Scatter plot of slow-decaying versus fast-decaying amplitudes demonstrates consistently larger slow-decaying responses. (l) Box plots of decay time constants for fast- and slow-decaying components showing similar kinetics. Nb. in one recording, only a slow (but no fast) component could be identified (see Methods for the statistical criterion distinguishing one- from two-component model fits). Consequently, this ‘unpaired’ slow decay is shown in red and has no corresponding paired value in the fast condition.

To corroborate the inference that the slow component of the ChI-evoked PSC was attributable to NGFI-driven engagement of extrasynaptic GABA_A_Rs, the DLS of mice expressing optimized FLP recombinase (FLPo) under control of the neuron-derived neurotrophic factor (NDNF) promoter ^27^ was injected with an AAV carrying a FLP-dependent Chrimson expression construct. NDNF is robustly expressed by NGFIs in both the striatum and cortex ^28^. In these experiments, iSPNs were identified by A2a-eGFP fluorescence and dSPNs by its absence (Fig. 3g, h). The NGFI-evoked PSCs in SPNs were typically greater than 100 pA and dominated by a slow current that was kinetically similar to the slow component deconstructed from ChI-evoked currents (Fig. 3i-l). The fast component of the NGFI-evoked PSCs could be a consequence of gap junction coupling to neighboring THIs ^25^.

### ChI engagement of GIs modulates SPN dendritic integration

To gain insight into how ChI control of GABAergic interneurons might shape SPN dendritic integration, a computational approach was taken. Our previous work has shown that the reversal potential for GABA_A_Rs in adult SPNs is near −60 mV and that ChI-evoked GABAergic input to ‘resting’ or down-state SPNs (~−85 mV) is depolarizing ^16^. Although spatiotemporally overlapping GABAergic and glutamatergic input to SPN dendrites can result in destructive interference ^29^, less spatially focused GABAergic depolarization of dendrites can enhance the ability of glutamatergic inputs to engage N-methyl-D-aspartate receptors (NMDARs). and drive dendritic spike generation in SPNs ^16^. To rigorously explore the spatial and temporal features of this interaction, computational models of an iSPN and a dSPN were generated. These models took advantage of iSPN and dSPN anatomical reconstructions that captured differences in their dendritic trees, as well as what is known about somatic and dendritic ion channel distributions shaping their physiology ^29–31^. Model parameters were tuned to recapitulate experimental differences in the response to somatic current injection (Fig. S1), as well as the ability of distal dendrites (>90 µm from the soma) to generate regenerative potentials in response to spatiotemporally convergent glutamatergic input ^32^.

To better understand the implications of the kinetic differences in the ChI-evoked synaptic responses mediated by THIs and NGFIs, the impact of fast and slow GABAergic inputs on an iSPN were studied separately. A cluster of glutamatergic synapses (11 neighboring spines) was placed on a distal dendrite that was capable of regenerative activity. To simulate the THI and NGFI circuits, either twelve fast, 1000 pS GABAergic synapses or 170 slow, 25 pS GABAergic receptors were distributed across the dendritic tree of an iSPN. To mimic slow activation of extrasynaptic receptors, GABA_A_Rs in NGFI circuits had an order of magnitude slower kinetics. The peak depolarizations produced by the underlying GABAergic responses at the dendritic site of glutamatergic activity were of similar magnitude (Fig. 4a-b). The timing of the glutamatergic input was varied relative to the distributed GABAergic input and the membrane potential at the dendritic site of glutamatergic input and the soma monitored. As shown previously ^16^, when the glutamatergic input trailed the fast GABAergic synaptic input, it was augmented by the depolarization produced by the opening of GABA_A_Rs. However, when the glutamatergic input preceded the GABAergic input, there was a narrow time window in which the spatially distributed GABA_A_R opening could modestly attenuate the glutamatergic depolarization. NMDAR-dependent dendritic spikes were not observed at any temporal interval in control simulations (Fig. 4c,d). To determine how the well described M_1_ mAChR (M_1_R) modulation of somatodendritic K^+^ channels in iSPNs might alter the interaction between GABAergic and glutamatergic inputs, the conductances of K_ir_2, K_v_4 and K_v_7 K^+^ channels were reduced by half (M_1_R activation) and the simulations re-run. In this state, which was intended to mimic the modulation resulting from tonic ChI activity, the ability of fast GABAergic synapses to augment the depolarization by focused glutamatergic input was dramatically enhanced (Fig. 4c,d). In this state, there was a temporal window of about 60 msec in which there was a several fold increase in the depolarization produced by the glutamatergic synapses. Dendritic spikes were evoked not only when the glutamatergic input trailed the GABAergic input, but also for a brief period when the timing was reversed (Fig. 4d). Varying the number of glutamatergic synapses revealed that the M_1_R modulation decreased the number of glutamatergic synapses necessary to evoke a dendritic spike with concomitant GABAergic input and increased the resulting somatic depolarization, resulting in somatic spikes with ~12 glutamatergic synapses (Fig. S2a-b).

**Figure 4.**
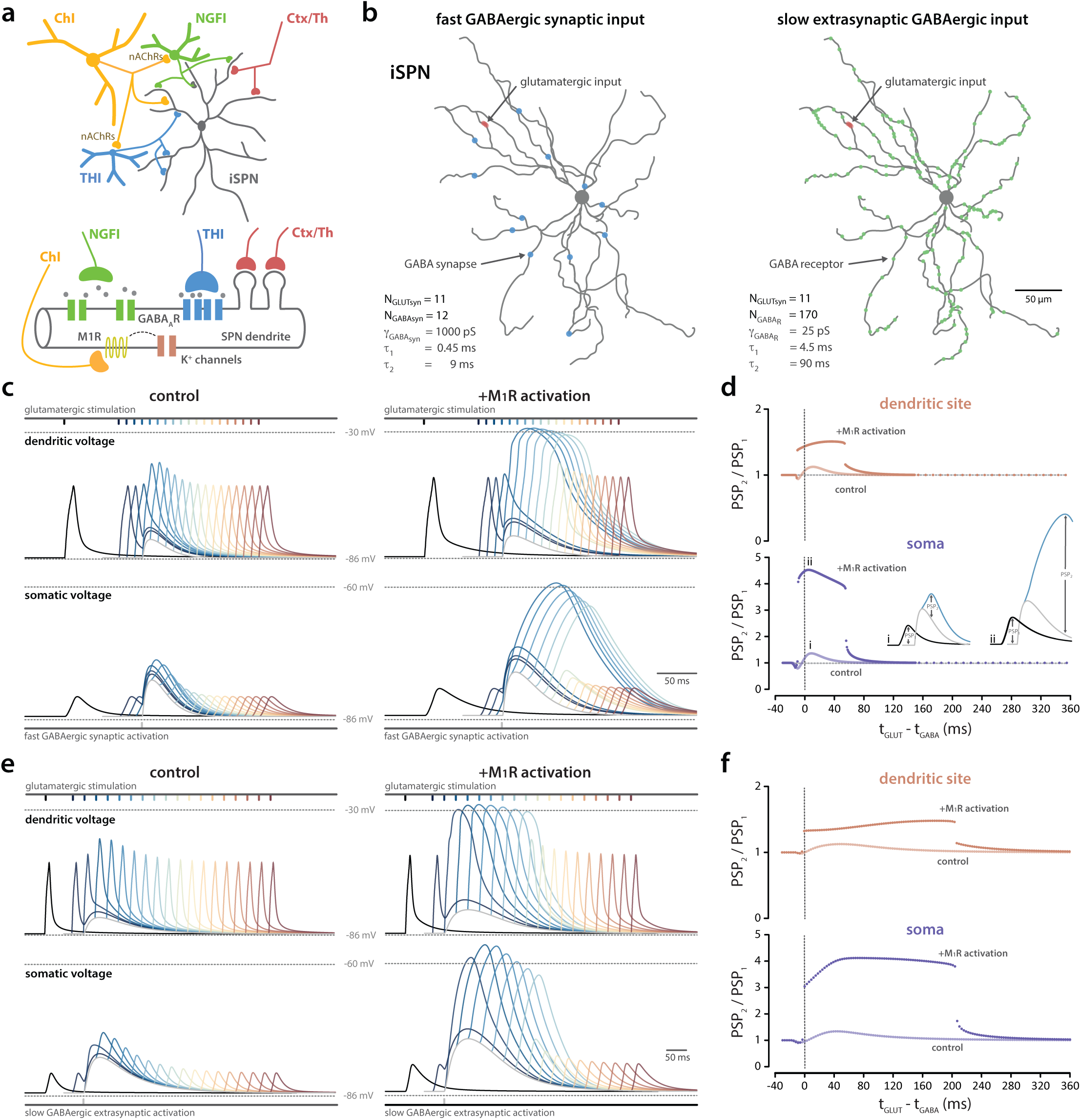
Effect of M_1_ mAChR activation on the temporal interaction between subthreshold glutamatergic and GABAergic inputs in a model iSPN. (a) Circuit diagram depicting cholinergic control of GABAergic microcircuits targeting SPNs and a schematic illustrating inputs onto a stretch of SPN dendrite. Corticothalamic (Ctx/Th) glutamatergic input onto dendritic spines provides fast excitatory drive. Cholinergic activation of NGFI (green) and THI (blue) neurons drives slow extrasynaptic GABAergic receptor activation and fast synaptic GABAergic input on the target SPN (black), respectively. (b) Reconstructed iSPN morphology showing dendritic sites of clustered glutamatergic activation (red) and fast synaptic GABAergic input (blue; left) and slow extrasynaptic GABAergic receptor activation (green; right). (c) Fast synaptic GABAergic input effect on glutamatergic synaptic potentials at dendritic and somatic sites in control (left) and following **M_1_** receptor activation (right). Black trace shows subthreshold glutamatergic excitation (11 neighboring spines at a single dendritic site). Gray trace shows phasic fast synaptic GABAergic activation (12 synapses each comprising 40 receptors with 25 pS peak conductance; Σg_GABA_ = 12 nS; PSP amplitude 8.0 mV at dendritic spike site). Color traces (blue to red) illustrate glutamatergic-GABAergic interactions across temporal offsets Δt = t_GLUT_ − t_GABA_ ranging from −30 to 150 ms; 10 ms intervals at dendritic site of activation and cell soma. (d) Summary of timing-dependent effects on the resultant PSP amplitude at dendritic (orange) and somatic (blue) sites, normalized to control, in the absence and presence of **M_1_** receptor activation at increased temporal offset fidelity. (e, f) As in (b, c) but for slow extrasynaptic GABAergic receptor activation (170 receptors, 25 pS; Σg_GABA_ = 4.25 nS; PSP = 7.3 mV at dendritic spike site) with Δt ranging from −30 to 480 ms; 30 ms intervals. Fast synaptic GABAergic input and slow extrasynaptic GABAergic receptor activation both augmented PSPs produced by clustered glutamatergic input, but, in both simulations, remained subthreshold for dendritic spike generation under control conditions. Following **M_1_** receptor modulation of K_ir_2, K_v_4 and K_v_7 K^+^ channels, supralinear NMDA receptor-dependent dendritic spikes were observed. Slow extrasynaptic GABAergic receptor activation extended the temporal window for this enhancement.

To assess the impact of the slow NGFI GABAergic currents, a cluster of glutamatergic synapses was placed at the same dendritic site as above. In the control state, the slow, NGFI-like GABAergic input led to a very modest attenuation of the glutamatergic input when it came first. However, when it trailed the GABAergic input, the glutamatergic depolarization was augmented as with the fast GABAergic synapses, but the time window for this augmentation was much longer, reflecting the slow decay of the GABAergic depolarization (Fig. 4e, f). As above, the M_1_R suppression of K_ir_2, K_v_4 and K_v_7 K^+^ channels dramatically increased the magnitude of the GABAergic enhancement of the glutamatergic input, creating a broad ‘coincidence’ window of roughly 200 msec (~3X that of the fast GABAergic input). Moreover, the M_1_R modulation reduced the number of glutamatergic synapses necessary to generate a dendritic spike with coincident GABAergic input and increased the resulting somatic depolarization and spiking (Fig. S2c, d).

To characterize how GABAergic input shapes the response of iSPNs to suprathreshold glutamatergic input, the number of dendritic sites receiving clustered glutamatergic input was increased to four with each site having a clustered glutamatergic input (comprising 14 neighboring spines). Fourteen neighboring spines were the minimum glutamatergic drive capable of generating a robust local dendritic spike. When all four clusters were activated coincidentally, three somatic action potentials were generated (not shown). To simulate THI and NGFI inputs, either forty fast, 1000 pS GABAergic synapses or 1600 slow, 25 pS synapses were distributed across the dendritic tree of an iSPN. Both yielded the same overall total conductance (Fig. 5a,b). As above, the timing of the glutamatergic input relative to a spatially distributed, fast GABAergic input was varied. As expected, as the GABAergic input came into temporal alignment with the glutamatergic input, dendritic spikes were truncated and the number of somatic spikes dropped or spike latency was increased (Fig. 5c, left). Somewhat surprisingly, this inhibitory effect of GABAergic input was essentially lost when the M_1_R modulation was simulated (Fig. 5c, right). Plotting the number of somatic spikes as a function of the time difference between the two inputs showed that there was roughly an 80 msec time window in which a trailing GABAergic input suppressed spike generation; M_1_R modulation blunted this effect and prevented spike suppression (Fig. 5d). Varying the number of GABAergic synapses, revealed that the GABAergic PSP amplitude and the effect on somatic impedance monotonically increased with their number; the threshold for the inhibition of somatic spiking was about 20 synapses. Again, the M_1_R modulation eliminated spike suppression, even with 50 activated GABAergic synapses(Fig. S3a-b).

**Figure 5.**
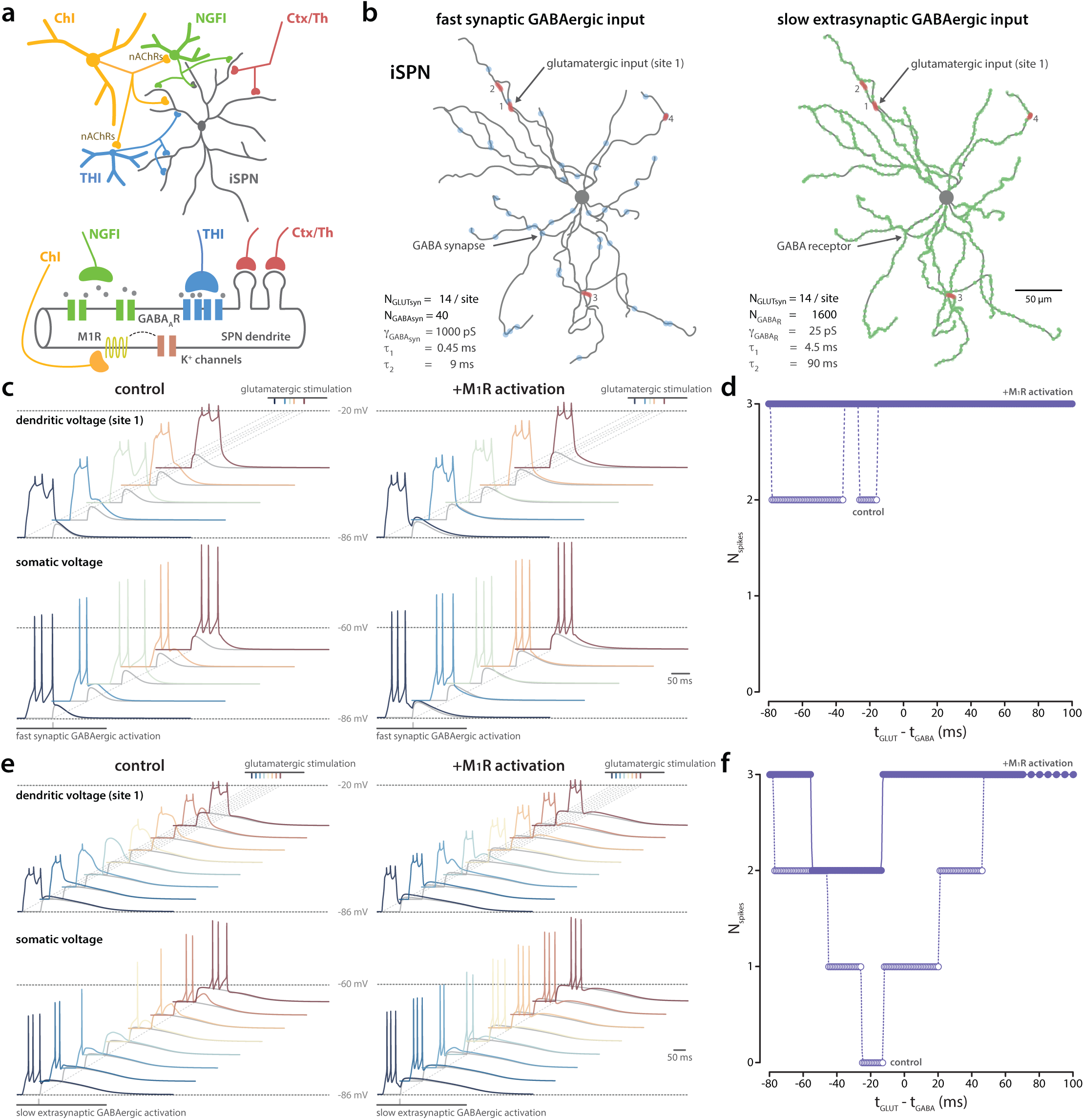
Effect of M_1_ mAChR activation on the temporal interaction between suprathreshold glutamatergic and GABAergic inputs in a model iSPN. (a) Circuit diagram depicting cholinergic control of GABAergic microcircuits targeting SPNs and a schematic illustrating inputs onto a stretch of SPN dendrite. (b) Reconstructed iSPN morphology showing four dendritic sites of clustered glutamatergic activation (red) and fast synaptic GABAergic input (blue; left) and slow extrasynaptic GABAergic receptor activation (green; right). (c) Effect of fast GABAergic activation on glutamatergic responses at dendritic and somatic sites in control (left) and following M_1_ receptor modulation of K_ir_2, K_v_4 and K_v_7 K^+^ channels (right). Black trace shows suprathreshold glutamatergic excitation (14 neighboring synapses activated at four independent dendritic sites; dendritic example taken from input site 1) and gray trace shows GABAergic activation (40 synapses each comprising 40 receptors with 25 pS peak conductance; Σg_GABA_ = 40 nS). Color traces illustrate interactions synaptic potentials at a dendritic site of clustered glutamatergic input (site 1) and at the cell soma for temporal offsets Δt = t_GLUT_ − t_GABA_ of −80, −50, −35, −20, 10 ms, respectively. (d) Summary of effects on somatic spike output across Δt in control and M_1_ receptor activation conditions. (e, f) As in (b, c) but for slow extrasynaptic GABAergic input (gray trace: 1600 receptors, 25 pS; Σg_GABA_ = 40 nS) with Δt = t_GLUT_ − t_GABA_ illustrated from −80 to 60 ms; 20 ms intervals. In both simulations, GABAergic activation reduced the total number of spikes generated at the soma. When repeated with concomitant M_1_R activation, the suppression of spike activity was attenuated helping to maintain somatic train output. The suppression of spiking and its relief by M_1_R activation was more pronounced with slow GABAergic activation.

Next, these simulations were repeated with distributed, small conductance, slow GABAergic synapses in an attempt to mimic the NGFI regulation of iSPNs. Not surprisingly, the slow GABAergic input more robustly inhibited dendritic and somatic spikes than the more transient, fast GABAergic synapses (Fig. 5e, left). Furthermore, the M_1_R modulation attenuated this inhibitory effect (Fig. 5e, right). Plots of somatic spike generation as a function of the temporal difference between inputs revealed that the window within which suppression took place was much broader than with the fast GABAergic input. Moreover, unlike the fast GABAergic input, inhibition of spiking was evident even when the GABAergic input preceded the glutamatergic input (*i.e*. t_GLUT_ − t_GABA_ < 0; Fig. 5f). As above, the amplitude of the GABAergic PSP increased monotonically with the number of synapses and somatic impedance fell in parallel. Furthermore, as the number of GABAergic synapses increased, the suppression of somatic spiking became more effective. Lastly, simulating the M_1_R modulation blunted this inhibition, with only a modest suppression of spiking at the greatest number of GABAergic synapses (Fig. S3c, d).

The dSPN simulations yielded very similar results for subthreshold and suprathreshold glutamatergic inputs (see Fig. S4a,b for simulation specifications). Distributed, fast GABAergic inputs boosted the response to subthreshold glutamatergic inputs at both the dendritic site, as well as the soma. The magnitude and time dependence of the interaction was nearly identical to that in iSPNs (Fig. S4c). What was profoundly different was the effect of mAChR modulation. Unlike iSPNs, M_1_Rs in dSPNs are only negatively coupled to K_v_7 K^+^ channels, which are engaged near spike threshold ^33,34^. M_4_Rs expressed by dSPNs are not known to directly modulate other dendritic or somatic ion channels involved in dendritic integration ^35,36^. As a consequence, mAChR signaling did not significantly modify the interaction between the fast GABAergic and glutamatergic inputs (Fig. S4c, d). The simulations of the slow, NGFI input to dSPNs also yielded results similar to those in iSPNs (see Fig. 6a-b for simulation specifications). The slow GABAergic input enhanced the response to trailing subthreshold glutamatergic inputs, but this enhancement was not significantly altered by simulating M_1_R stimulation (Fig. 6c, d). When the glutamatergic input was expanded to create a suprathreshold synaptic event, the slow GABAergic input significantly inhibited spike generation (Fig. 6e, f). Interestingly, this inhibition occurred when the GABAergic input preceded or followed the glutamatergic input, but was most profound when the slow GABAergic PSP slightly trailed the glutamatergic input (5-20 msec) (Fig. 6f). The M_1_R modulation only modestly attenuated this inhibition (Fig. 6e, f).

**Figure 6.**
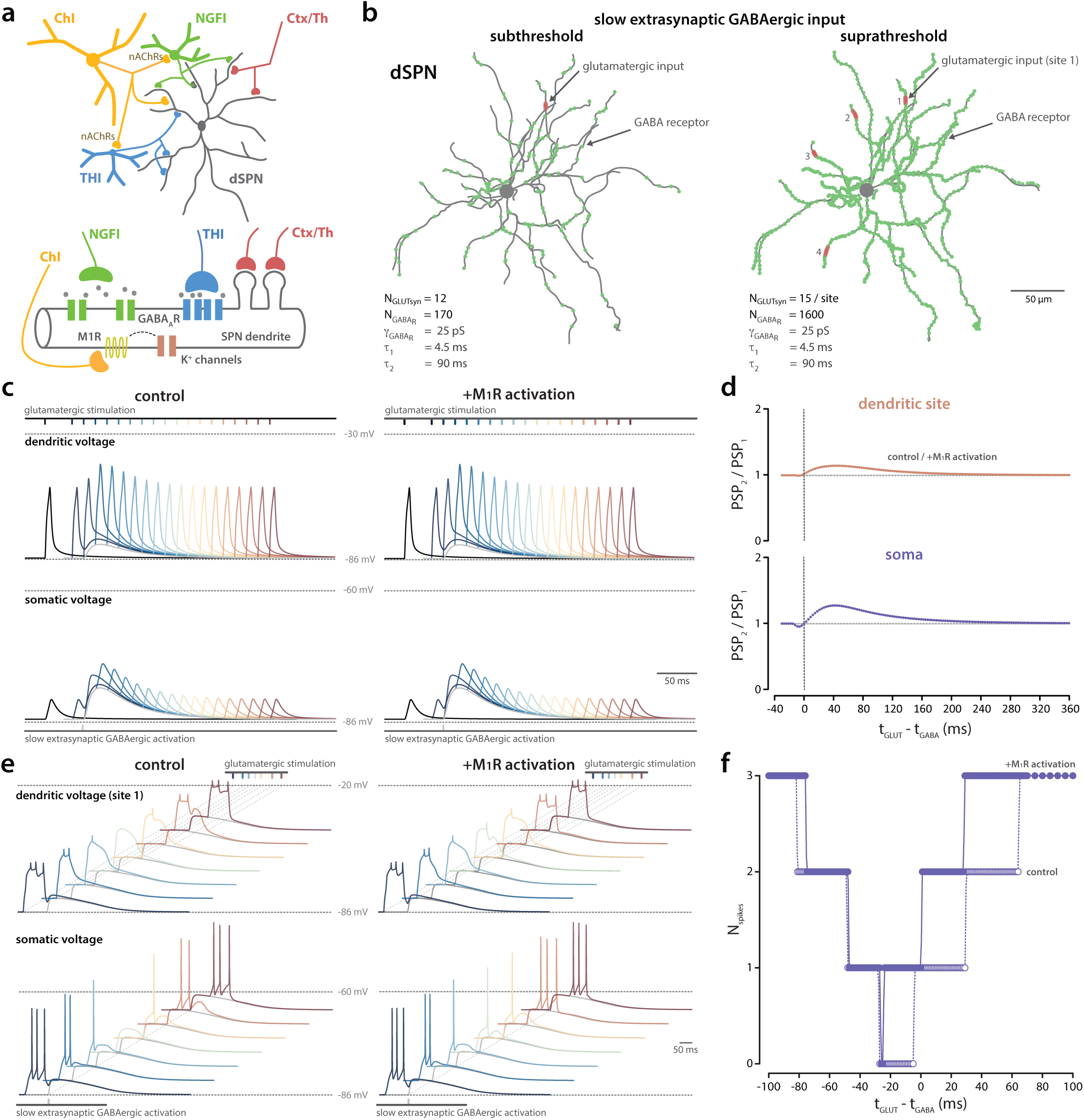
Effect of M_1_ mAChR activation on the interaction between glutamatergic input and slow extrasynaptic GABAergic responses in a model dSPN. (a) Circuit diagram depicting cholinergic control of GABAergic microcircuits targeting SPNs and a schematic illustrating inputs onto a stretch of SPN dendrite. (b) Reconstructed dSPN morphology showing subthreshold (1 site; left) and suprathreshold (4 sites; right) of clustered glutamatergic activation (red) and slow (green; right) GABAergic inputs. **(c)** Slow extrasynaptic GABAergic activation effect on glutamatergic synaptic potentials at dendritic and somatic sites in control (left) and following M_1_ receptor modulation of K_v_7 channels (right). Black trace: subthreshold glutamatergic excitation (12 inputs). Gray trace: slow GABAergic activation (170 receptors, 25 pS; Σg_GABA_ = 4.25 nS; PSP = 6.2 mV at dendritic spike site). Colored traces (blue to red): glutamatergic-GABAergic interaction at temporal offsets Δt = t_GLUT_ − t_GABA_ is illustrated from −30 to 480 ms; 30 ms intervals. (d) Summary of timing dependent effects on PSP amplitude. No effect of M_1_ receptor activation was observed. (e, f) As in (c, d) but for suprathreshold glutamatergic input (15 inputs per site; 4 sites). Gray trace: slow extrasynaptic GABAergic activation (1600 receptors, 25 pS; Σg_GABA_ = 40 nS). Colored traces (blue to red): glutamatergic-GABAergic interaction at temporal offsets Δt = t_GLUT_ − t_GABA_ of −100, −60, −40, −20, 0, +40 and +80, respectively. Only a modest effect of M_1_ receptor activation on spike output was observed.

Taken together, these simulations suggest that ChI-driven activation of THIs and NGFIs enhances temporally coincident glutamatergic input to iSPNs and dSPNs when these inputs are subthreshold for somatic spike generation. The coincidence ‘window’ for this effect is narrow for fast THI GABAergic input and several fold broader for the slow NGFI input. In iSPNs, but not dSPNs, this enhancement can be dramatically increased by M_1_R-mediated suppression of K_ir_2, K_v_4 and K_v_7 K^+^ channels. When SPNs are driven to spike by stronger, spatially distributed glutamatergic input, temporally aligned THI and NGFI GABAergic inputs are capable of suppressing dendritic and somatic spike generation, but in iSPNs this inhibitory effect is significantly attenuated by concomitant M_1_R signaling and negative modulation of K^+^ channels.

### In models of early-stage PD, ChI release of ACh was elevated

As mentioned above, striatal cholinergic signaling increases in PD ^19^. Indeed, in mouse models of late-stage PD created by 6-OHDA lesioning of the dopaminergic projection to the striatum, ChI-evoked ACh release in the DLS is elevated ^20^. What is less clear is whether the loss of striatal dopaminergic signaling in the absence of the broader network pathophysiology accompanying the parkinsonian state is sufficient to increase ChI-evoked ACh release. In principle, this should be the case as inhibitory signaling through presynaptic D_2_Rs should be lost. To test this hypothesis, ChI-evoked ACh release in the DLS of prodromal MCI-Park mice was assessed. Specifically, an AAV carrying an expression construct for the GRAB_ACh3.0_ sensor was stereotaxically injected into the DLS of P25 MCI-Park mice and litter-mate controls ^37^. Two weeks later at P40, mice were sacrificed and brain slices prepared. The DLS was electrically stimulated and two photon laser scanning microscopy (2PLSM) used to assess evoked changes in probe fluorescence (Fig. 7a). The evoked fluorescence response rose rapidly and fell back to baseline within a few seconds in both control and prodromal MCI-Park striata (Fig. 7b). Although similar in kinetics, the amplitude of the ACh transient was significantly larger in prodromal MCI-Park striata (Fig. 7c); these results suggest that, as in models of late-stage PD ^20^, ChI-evoked ACh release is dis-inhibited in the prodromal striatum.

**Figure 7.**
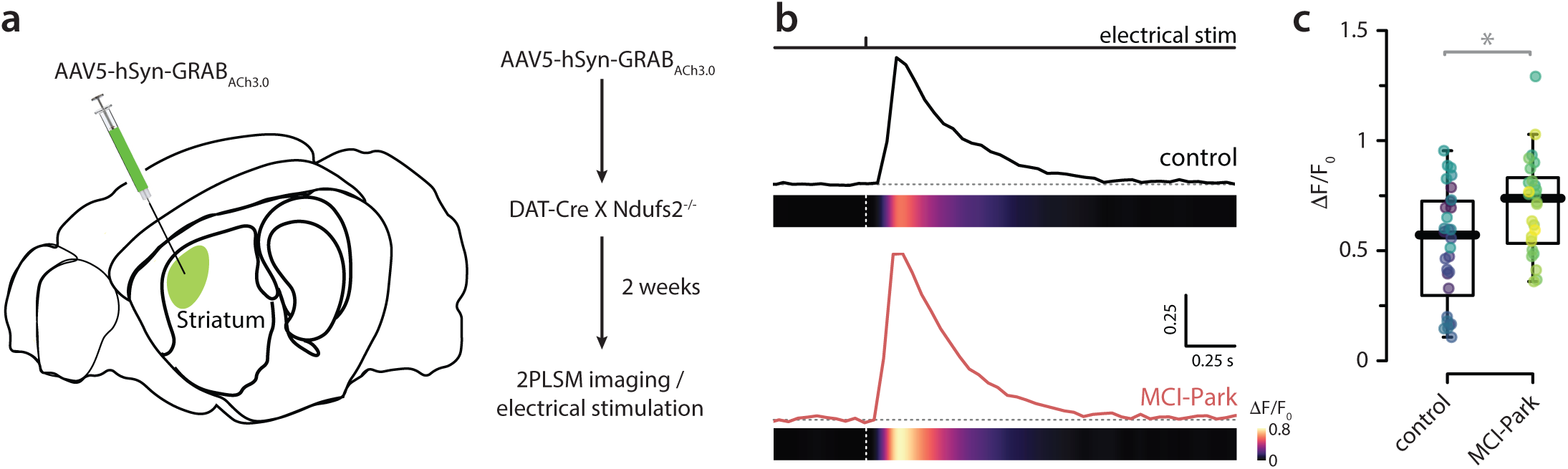
ChI-evoked acetylcholine release was elevated in MCI-Park mice. (a) Schematic of viral injection of GRAB-ACh3.0 into the DLS of DAT-Cre x Ndufs2^−/−^ mice and experimental timeline. (b) Representative traces of ChI-evoked ACh release in response to electrical stimulation (300 µA, 1 ms) in control and MCI-Park mice. (c) Box plots of normalized ACh responses showing elevated release in MCI-Park mice (robust linear mixed-effects model, cluster bootstrap *p* = 0.04620, N_boot_ = 9,999).

### In PD models, ChI activation of GI circuits coupled to SPNs was disrupted

Given the enhanced release of ACh by ChIs in both prodromal and parkinsonian states, our working hypothesis was that ChI coupling to THIs and NGFIs would be strengthened, leading to enhanced GABAergic input to SPNs. To test this hypothesis, MCI-Park mice were crossed with mice expressing Flp recombinase under the control of choline acetyl transferase promoter (*ChAT-Flp*). In P25 triple crosses (*DAT-Cre*^+/−^:: *Ndufs2^−/−^* :: *ChAT-Flp*) and age-matched controls (*DAT-Cre*^+/−^:: *ChAT-Flp*), an AAV carrying a Flp-dependent Chrimson expression construct was stereotaxically injected into the DLS. Two weeks later, ChI coupling to SPNs was evaluated using optogenetics in conjunction with whole cell voltage clamp electrophysiology in *ex vivo* brain slices. In these prodromal MCI-Park mice, striatal expression of tyrosine hydroxylase (TH) is virtually gone (Fig. 8a) and evoked dopamine release dramatically down-regulated ^23^. Unexpectedly, optogenetic stimulation of ChIs in tissue from prodromal MCI-Park mice produced significantly smaller PSCs (both peak and total charge transfer) in SPNs than the same stimulation did in controls (Fig. 8b-d). Although the kinetics of the PSCs was unaltered, the amplitude of both fast and slow components of the PSC were significantly smaller in prodromal MCI-Park SPNs (Fig. 8e, f).

**Figure 8.**
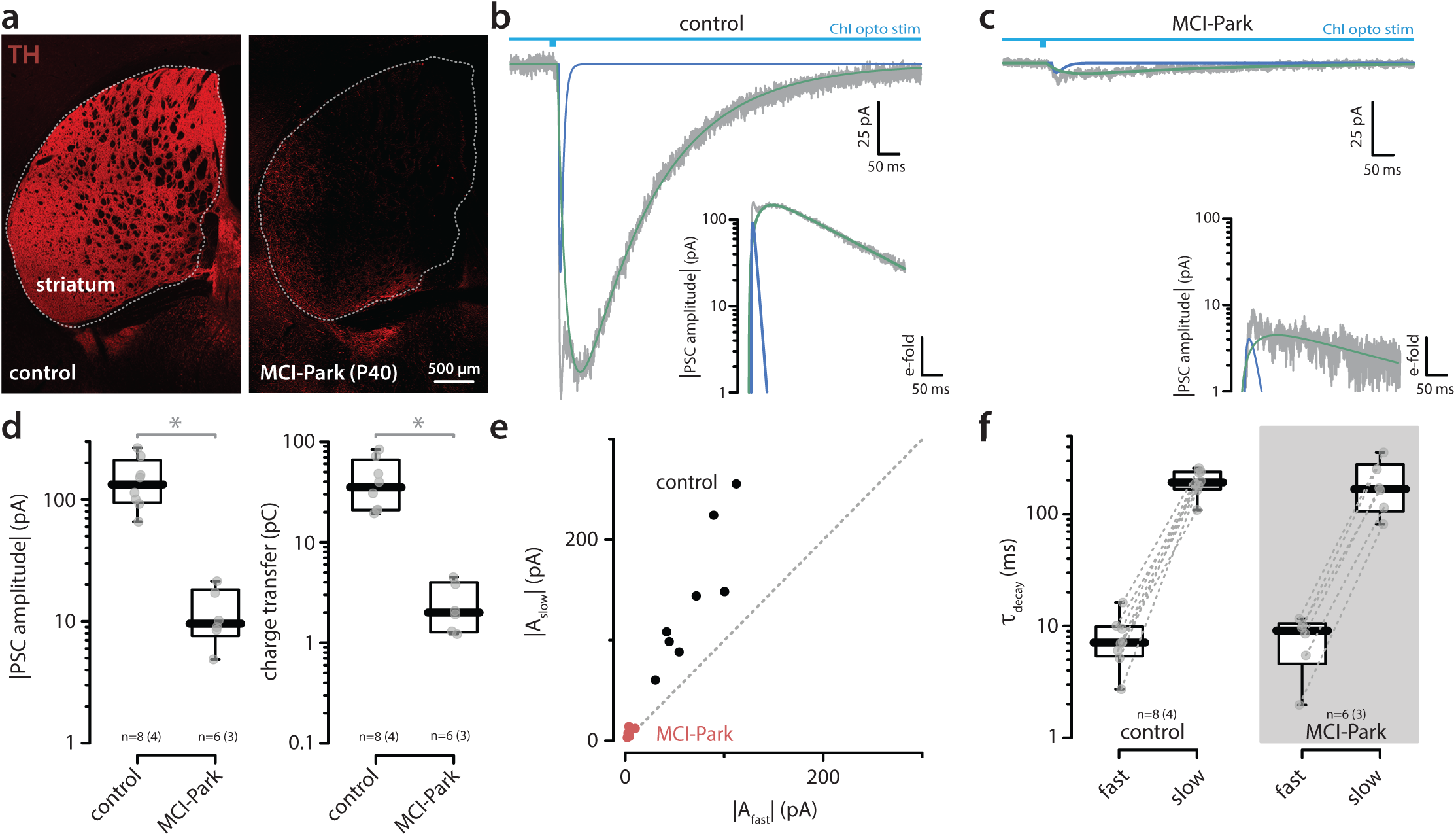
ChI-evoked GABAergic PSCs were attenuated in MCI-Park mice. (a) Confocal images of striatal sections stained with TH antibody showing dopamine loss at P40 in MCI-Park mice. Representative PSCs recorded from SPNs in control (b) and MCI-Park (c) mice in response to activation of ChIs (whole-field LED, 5 ms), showing fast- and slow-decaying gabazine-sensitive components. Insets, semi-logarithmic scale. (d) Semi-log box plots of PSC amplitudes and charge transfer in control and MCI-Park conditions showing reduced responses in MCI-Park mice (Wilcoxon rank-sum test: amplitude, W = 0, p = 0.00067; charge transfer, W = 48, p = 0.00067). (e) Scatter plot of slow-decaying versus fast-decaying amplitudes. (f) Semi-log box plots of decay time constants in control and MCI-Park conditions showing no change in kinetics (Wilcoxon rank-sum test, control vs MCI-Park: fast, W = 15, p = 0.56477; slow, W = 29, p = 0.57276). Note in one recording in the MCI-Park group, only a slow (but no fast) component could be identified (see Methods for the statistical criterion distinguishing one- from two-component model fits). Consequently, this ‘unpaired’ slow decay is shown in red and has no corresponding paired value in the fast condition. Displayed n values are adjusted accordingly.

To determine whether this uncoupling extended to a model of late-stage PD, the unilateral 6-hydroxy dopamine (6-OHDA) lesion model was studied. The DLS of ChAT-Cre:: *Drd1a^tdTomato^* or ChAT-Cre:: *Drd2^e^*^GFP^ mice was stereotaxically injected with an AAV carrying a Cre-dependent expression construct encoding Chronos (Fig. 9a). In parallel, 6-OHDA was injected into the medial forebrain bundle (MFB). Four weeks later, the extent of the lesion was assessed using a forelimb asymmetry test ^38^. As expected, mice with strong asymmetry scores (<30% contralateral paw usage) had a profound loss of striatal TH immunoreactivity (Fig. 9b). In agreement with the experiments in MCI-Park mice, optogenetic stimulation of ChIs in *ex vivo* brain slices from lesioned mice produced much smaller GABAergic PSCs in both iSPNs and dSPNs (Fig. 9c-g). Both the fast and slow components of the PSC were smaller in 6-OHDA lesioned striata (Fig. 9h), but the kinetics of the responses were not discernibly different (Fig. 9i). Similar results were obtained in independent experiments (Fig. S5).

**Figure 9.**
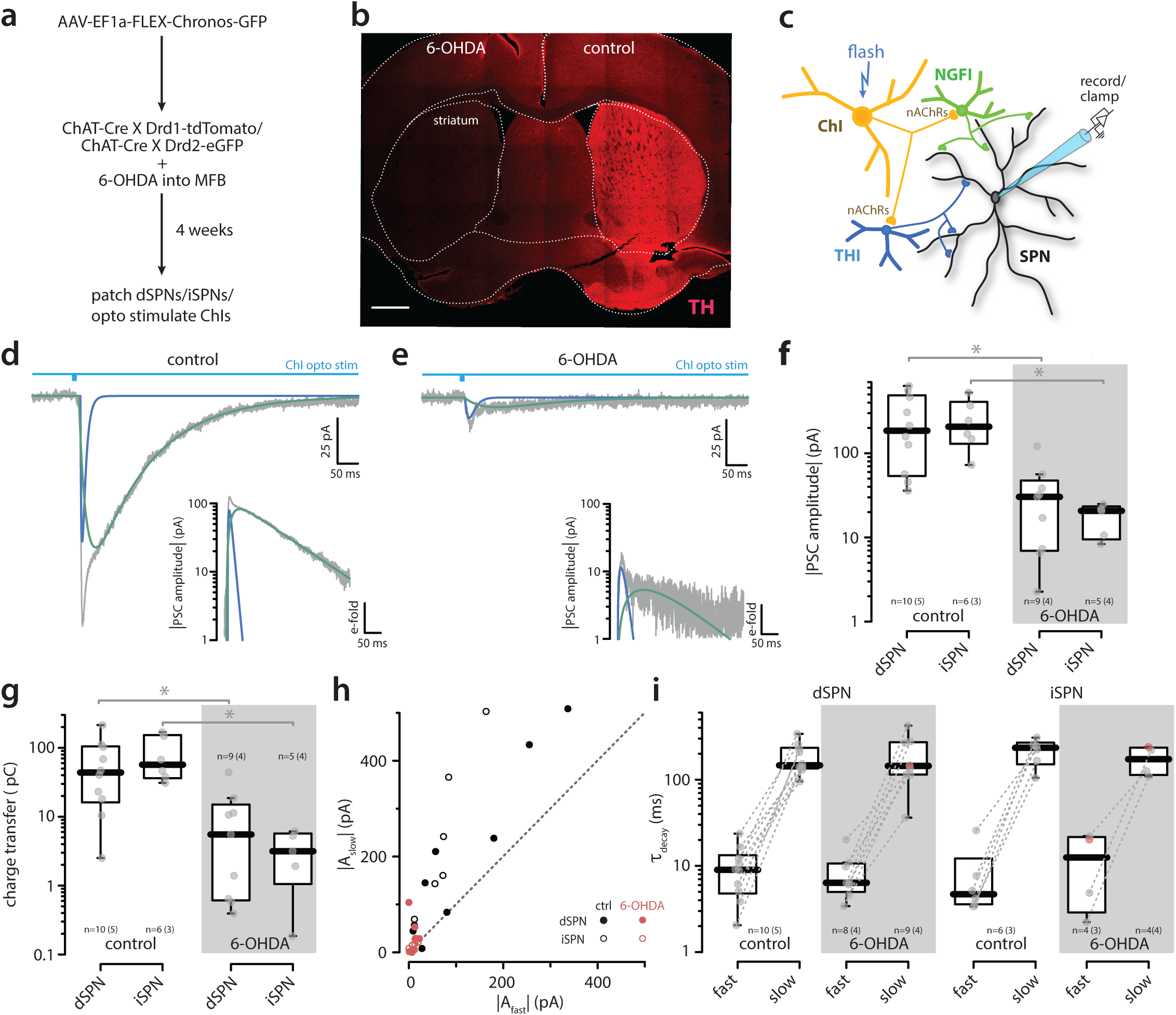
ChI-evoked GABAergic input in SPNs was reduced following 6-OHDA lesion. (a) Schematic of opsin injection into the DLS together with 6-OHDA injection into the medial forebrain bundle in ChAT-Cre x D_1_-tdTomato mice, and experimental timeline. (b) Striatal coronal section stained with TH antibody four weeks after 6-OHDA injection. Representative PSCs recorded from SPNs in control (c) and 6-OHDA (d) conditions in response to activation of ChIs (whole-field LED, 5 ms), showing fast- and slow-decaying gabazine-sensitive components. Insets, semi-logarithmic scale. (e) Semi-log box plots of gabazine-sensitive PSC amplitudes in dSPNs (control: n = 10, 5 animals; 6-OHDA: n = 9, 3 animals) and iSPNs (control: n = 6, 4 animals; 6-OHDA: n = 5, 4 animals) showing reduced responses after 6-OHDA (Wilcoxon rank-sum test, control vs 6-OHDA: dSPN, W = 6, *p* = 0.00260; iSPN, W = 0, *p* = 0.01299). (f) Semi-log box plots of total charge transfer for the same cells showing reduced responses after 6-OHDA. (g) Scatter plots of slow-decaying versus fast-decaying amplitudes in control (black) and 6-OHDA (red) conditions (Wilcoxon rank-sum test, control vs 6-OHDA: dSPN, W = 77, p = 0.02286; iSPN, W = 30, p = 0.01732). Open and closed symbols are used for dSPN and iSPN, respectively. (h) Semi-log box plots of fast and slow decay time constants in dSPNs and iSPNs showing no change in kinetics (Wilcoxon rank-sum test, control vs 6-OHDA: dSPN fast, W = 44, *p* = 1; iSPN fast, W = 11, *p* = 1; dSPN slow, W = 48, *p* = 1; iSPN slow, W = 17, *p* = 1). Note, in some recordings, only one component (fast or slow) was identified (see Methods for the statistical criterion). These ‘unpaired’ decays are shown in red without a corresponding value in the other condition. Displayed n values are adjusted accordingly.

### Coupling of GIs to SPNs was not altered in PD models

To determine whether dopamine depletion altered the coupling of THIs and NGFIs with SPNs, this linkage was interrogated using optogenetic approaches. First, mice expressing Cre recombinase under control of the neuropeptide Y (NPY) promoter were used to examine the coupling of GABAergic low threshold spiking interneurons (LTSIs) and NGFIs with SPNs. NPY-Cre mice were crossed with either *Drd1a^tdTomato^* or *Drd2^e^*^GFP^ transgenic mouse lines. An AAV carrying Cre-dependent Chronos construct was injected into the DLS of these mice on the same day as 6-OHDA was injected into the medial forebrain bundle (MFB) (Fig. 10a). Four weeks later, mice were sacrificed and brain slice prepared for study using the approaches outlined above; as expected, NPY and SPNs were readily identified in these slices (Fig. 10b). Somewhat surprisingly, the PSCs evoked in iSPNs and dSPNs by optogenetic activation of LTSIs and NGFIs were unaltered by the 6-OHDA lesion. Both the fast component, which was largely attributable to LTSIs, and the slow component, which was NGFI-dependent, were similar in amplitude and charge transfer (Fig. 10c-f). As visualized in a scatter plot (Fig. 10g), the distribution of fast and slow PSC amplitudes in iSPNs and dSPNs from control and lesioned mice was overlapping, indicating that there was no change in this synaptic linkage. In agreement with this conclusion, the kinetics of the fast and slow components of the PSC were unaltered by lesioning (Fig. 10h).

**Figure 10.**
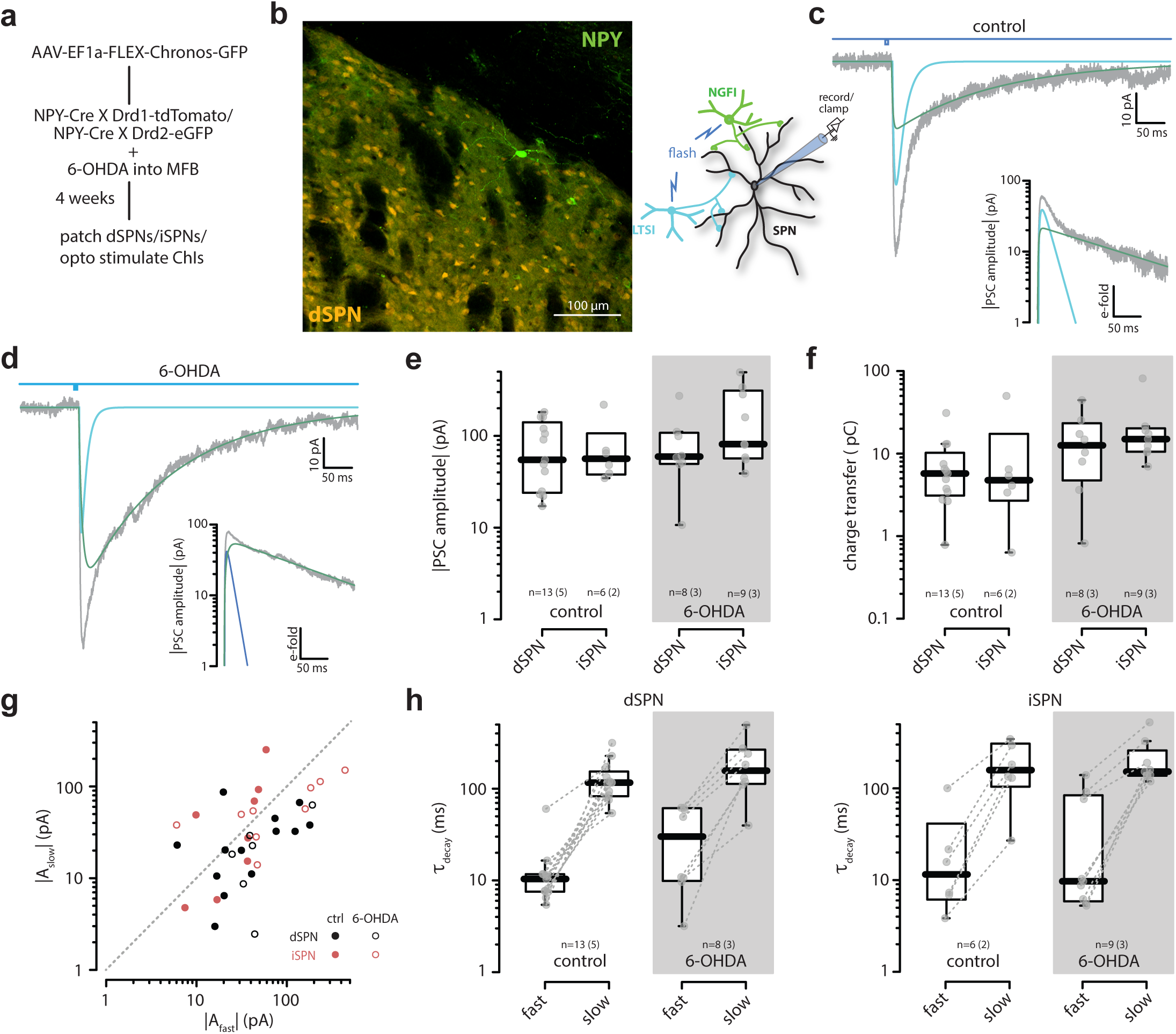
NPY-evoked GABAergic input in SPNs remained intact following 6-OHDA lesion. (a) Experimental timeline including viral injection and 6-OHDA lesion. (b) Confocal image of a coronal striatal section from NPY-Cre x D_1_-tdTomato mice showing Chronos expression in NPY interneurons (green) and tdTomato in dSPNs (orange). Representative gabazine-sensitive PSCs recorded from SPNs in control (c) and 6-OHDA (d) conditions in response to activation of NPY interneurons (whole-field LED, 5 ms), showing fast- and slow-decaying components. Insets, semi-logarithmic scale. (e) Semi-log box plots of gabazine-sensitive PSC amplitudes in dSPNs (control: n = 13, 5 animals; 6-OHDA: n = 8, 3 animals) and iSPNs (control: n = 6, 2 animals; 6-OHDA: n = 9, 3 animals) showing no change after 6-OHDA (Wilcoxon rank-sum test, control vs 6-OHDA: dSPN, W = 55, *p* = 1; iSPN, W = 40, *p* = 0.57862). (f) Semi-log box plots of total charge transfer showing no change after 6-OHDA (Wilcoxon rank-sum test, control vs 6-OHDA: dSPN, W = 30, *p* = 0.36376; iSPN, W = 8, *p* = 0.10230). (g) Semi-log scatter plots of slow-decaying versus fast-decaying amplitudes in control (black) and 6-OHDA (red) conditions. Open and closed symbols are used for dSPN and iSPN, respectively. (h) Semi-log box plots of fast and slow decay time constants in dSPNs and iSPNs showing no change in kinetics (Wilcoxon rank-sum test, control vs 6-OHDA: dSPN fast, W = 34, p = 0.84104; iSPN fast, W = 26, *p* = 1; dSPN slow, W = 33, *p* = 0.73867; iSPN slow, W = 23, *p* = 1).

### The attenuation in ChI coupling was attributable to suppression of GI nAChRs

If the synaptic coupling of THIs and NGFIs with SPNs was unaltered by dopamine depletion, then the most likely explanation for the broader circuit deficit was that the coupling between ChIs and interneurons was disrupted. To test this hypothesis, the expression of mRNA coding for the nAChR β2 subunit was examined using RNAscope. Previous work has shown that β2-containing nAChRs are present at these synapses ^25^. To colocalize the β2 subunit probe with GIs, TH and NDNF mRNA probes were used to identify THIs and NGFIs. Tissue from control and 6-OHDA lesioned mice (as described above) was examined using these three RNAscope probes (Fig. 11a). Grain counts for the β2 subunit probe were modestly, but significantly, reduced in tissue from 6-OHDA lesioned mice (Fig. 11b). The reduction in β2 subunit expression was similar in NDNF and TH interneurons (Fig. S6).

**Figure 11.**
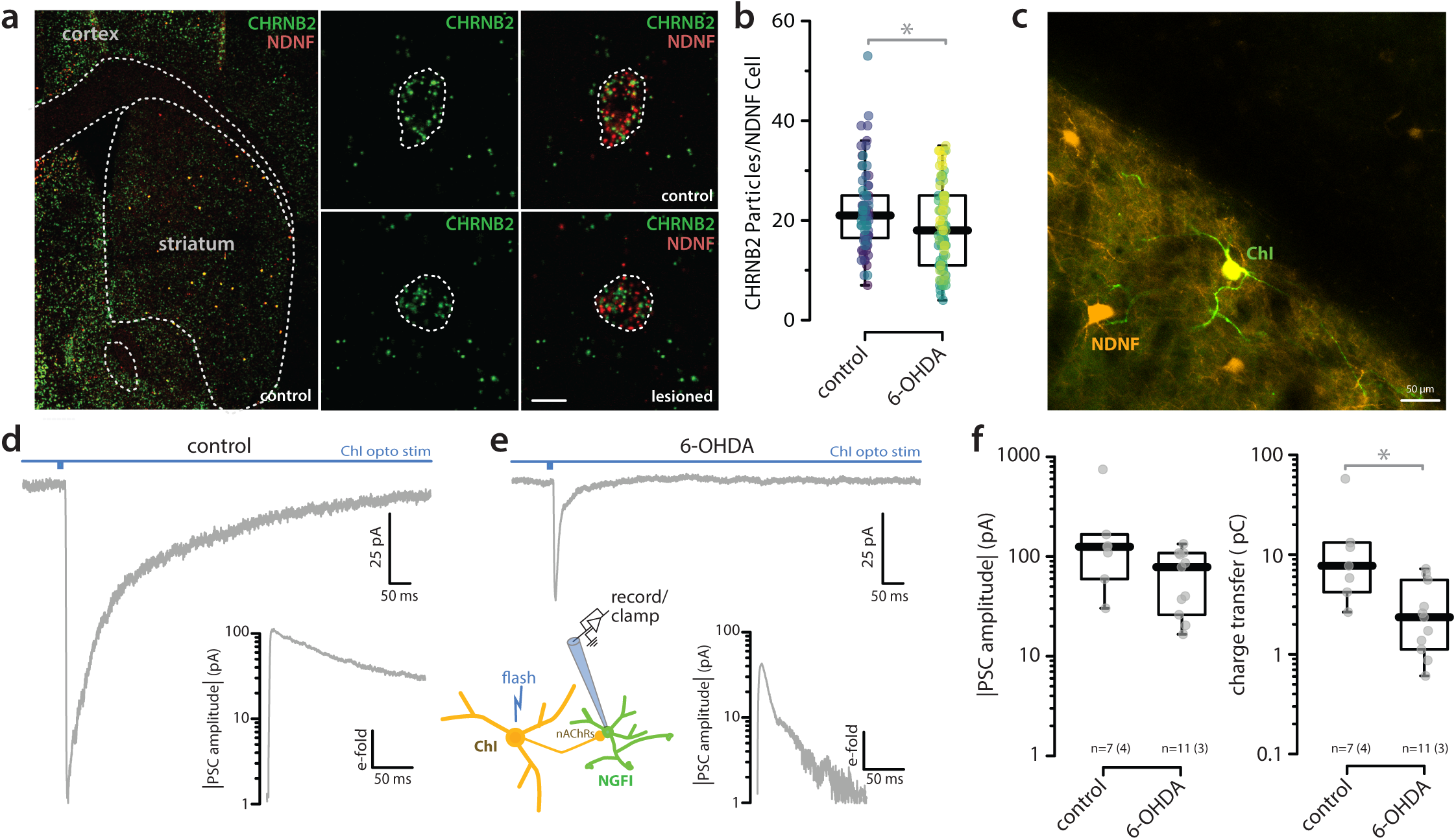
ChI-evoked input to NGF interneurons was reduced following 6-OHDA lesion. (a) In situ hybridization showing CHRNB2 (β2 subunit) mRNA colocalized with NDNF in NGF interneurons in control and 6-OHDA conditions. (b) Box plots of β2 subunit-containing nAChR expression in NGF interneurons showing reduced expression after 6-OHDA (robust linear mixed-effects model, Nboot = 1,999, *p* = 0.00400). (c) Confocal image of a striatal section from ChAT-Cre x NDNF-Flp mice showing Chronos expression in ChIs (green) and mCherry expression in NGF interneurons (orange). Representative PSCs recorded from NGF interneurons in response to activation of ChIs (whole-field LED, 5 ms) under control (d) and 6-OHDA (e) conditions. (f) Semi-log box plots of PSC amplitudes showing no change, and charge transfer showing reduced responses after 6-OHDA (n = 7 cells, 4 animals; n = 11 cells, 3 animals) (Wilcoxon rank-sum test, control vs 6-OHDA: amplitude, W = 17, *p* = 0.05556; charge transfer, W = 68, *p* = 0.00591).

To test the functional connectivity of ChIs with NGFIs, mice expressing Cre recombinase under ChAT promoter (ChAT-Cre) were crossed with mice expressing Flp recombinase under control of the NDNF promoter (NDNF-Flp). An AAV carrying Cre-dependent Chronos construct was stereotaxically injected into the DLS of these mice along with an AAV carrying a Flp-dependent mCherry expression construct (Fig. 11c). A cohort of these mice were lesioned with 6-OHDA as described above. Four weeks later, control and lesioned mice were sacrificed and brain slices were prepared. NDNF-expressing NGFIs were visually identified and subjected to whole cell voltage clamp. In slices from control mice, optogenetic activation of ChIs elicited robust PSCs. However, in slices from 6-OHDA-lesioned mice, the ChI-evoked responses in NDNF/NGFIs were smaller. Although the peak current evoked by stimulation of ChIs was not significantly reduced, the total charge transfer was, as the currents decayed more rapidly (Fig. 11d-f). Taken together, these experiments suggest that the elevation in cholinergic signaling produced by dopamine depletion down-regulates THI and NGFI nAChRs, reducing the ability of ChIs to drive spiking.

## Discussion

There are three main conclusions that can be drawn from our experiments. First, in agreement with previous studies, our experiments demonstrate that ChIs are robustly coupled through GIs to both iSPNs and dSPNs. Furthermore, the strength of this disynaptic couple was similar in iSPNs and dSPNs, with the slower NGFI-mediated component being attributable in part to extrasynaptic GABA_A_Rs. Second, modeling studies suggest that this ChI-anchored circuit can create a temporal window in which extra-striatal glutamatergic inputs to SPNs are bidirectionally modulated. This ‘coincidence’ modulation could allow set-shifting events to trigger alterations in SPN ensemble architecture and movement. Third, in models of both prodromal and parkinsonian states, the coupling of ChIs to iSPNs and dSPNs was dramatically down-regulated, despite disinhibition of ACh release. The attenuated coupling was attributable in part to a down-regulation in surface expression of nAChRs in THIs and NGFIs, not to an alteration in GABAergic interneuron coupling to SPNs. This loss of ChI coupling could contribute to early deficits in set shifting, reversal learning and movement sequencing in prodromal, as well as in clinically defined PD patients.

This study confirms ChIs are robustly, di-synaptically coupled to both iSPNs and dSPNs. As shown previously ^15^, optogenetic activation of ChIs evoked a kinetically complex GABA_A_R-mediated synaptic response in SPNs consisting of fast and slow components. These two components have been attributed to nAChR-dependent activation of THIs and NGFIs ^15,25^. To quantitatively assess the contribution of these two circuit elements, mathematical methods were used to deconstruct evoked currents into slow and fast components. The decay time constants fitted to fast and slow currents differed by approximately an order of magnitude (facilitating their separation) in both iSPNs and dSPNs. Although the absolute amplitude of these components varied from cell-to-cell, the relative amplitudes of the two components did not significantly differ between iSPNs and dSPNs, showing that both cell types were robustly coupled to ChIs through this interneuron circuit.

The slower component of the ChI-evoked current has been attributed to activation of NGFIs and engagement of extrasynaptic GABA_A_Rs ^15,25^, as in other brain regions ^39^. Indeed, selective optogenetic activation of NGFIs in an NDNF-Flpo mouse line yielded synaptic currents in SPNs dominated by slow currents similar to those evoked by ChI stimulation. However, there was also a fast current component evoked by optogenetic stimulation of NGFIs. This fast current could stem either from activation of neighboring THIs coupled by gap junctions to NGFIs or a synaptic component of the NGFI innervation of SPNs ^15,25^. As pharmacological tools for manipulating gap junctions are problematic, additional studies using genetic strategies to disrupt connexin expression in NGFIs will be necessary to sort out the mechanisms underlying the fast component. Consistent with the inference that the slow kinetics of this current were attributable to extrasynaptic receptors, knocking down GABA_A_R δ subunits significantly reduced the amplitude of the slow component of the ChI-evoked response, as well as that evoked by NGFI stimulation. The retention of a substantial slow current after δ subunit knockdown could reflect the incompleteness of knockdown. Alternatively, it could reflect the participation of synaptic GABA_A_Rs that are physically remote from the NGFI release sites and are engaged only as GABA diffuses to them ^40^.

The computational studies presented here suggest that the ChI-anchored interneuron circuit regulates SPN dendritic integration. To gain a better perspective on the potential functional significance of the interneuronal circuitry, simulations were performed using anatomically accurate iSPN and dSPN models invested with ion channels known to contribute to their physiology ^16,29,41^. These studies focused on how dendritic GABAergic synapses, like those arising from THIs and NGFIs, might shape the responses of SPNs to glutamatergic input, particularly to distal dendrites that are capable of generating NMDAR-dependent spikes ^32^. In agreement with previous experimental and computational work ^16,42,43^, these simulations demonstrated that activation of dendritic GABA_A_Rs could serve as a state-dependent modulator of glutamatergic input: enhancing appropriately timed subthreshold glutamatergic inputs when SPNs were quiescent and suppressing them when the glutamatergic inputs were suprathreshold. In ‘resting’ SPNs, where the membrane potential is near the K^+^ equilibrium potential (~-85 mV), engagement of GABA_A_Rs (that reverse near −60 mV) depolarizes dendrites ^16,17^. When GABAergic input was spatially distributed, the impact on glutamatergic synaptic currents was mediated primarily by membrane depolarization, which brought the synaptic potential nearer that needed for relief of Mg^2+^ block of NMDARs and dendritic spike generation ^32^. The enhancement occurred primarily when the glutamatergic input trailed the GABA_A_R input (and receptors had closed), creating a temporal window in which coincident ChI and glutamatergic synaptic activity synergized. The breadth of this window was roughly three times longer when mediated by the nominally extrasynaptic, slow GABA_A_R input than that created by the fast GABAergic input.

When the dendritic glutamatergic inputs to SPNs were suprathreshold, spatially distributed GABAergic inputs, regardless of whether they were fast or slow, effectively reduced the glutamatergic depolarization and spiking. In these simulations, spatiotemporally convergent glutamatergic synapses were distributed across several locations in dendritic tree, each of sufficient magnitude to produce a local NMDAR-dependent spike and collectively sufficient to generate somatic spiking. Although qualitatively similar, the slow, nominally extrasynaptic GABAergic input was much more effective in suppressing somatic spike generation than was the fast, transient GABAergic input. This difference was largely attributable to the time course of the slower GABAergic PSP and its alignment with the slow dendritic spike produced by glutamatergic input.

ChIs also modulate the excitability of SPNs through postsynaptic mAChRs. In iSPNs, this signaling is mediated by G_q_-coupled M_1_Rs that suppress the opening of somatodendritic K^+^ channels ^33^. Interestingly, mimicking this postsynaptic modulation by reducing the density of K_v_4, K_v_7 and K_ir_2 K^+^ channels dramatically increased both the magnitude and duration of the GABA_A_R-mediated enhancement of subthreshold glutamatergic input and blunted the inhibitory effects on suprathreshold, multi-site input. This was true for both fast and slow GABAergic signaling, but understandably larger for the slow input. Thus, there clearly was a synergy between the nAChR- and mAChR-mediated regulation of quiescent iSPNs by ChIs, with both promoting the responsiveness to glutamatergic synapses and spiking. Interestingly, given that ACh acts presynaptically to inhibit intratelencephalic (IT) corticostriatal synapses and enhance those of pyramidal tract neurons ^14,35,44^, the coordinated effects of ChIs should preferentially strengthen the coupling of iSPNs to movement-related PT activity, perhaps serving to ‘pause’ ongoing movement.

In contrast, the modulation of dSPNs by ChIs is largely mediated by G_i/o_-coupled M_4_Rs ^36^ that are not discernibly coupled to dendritic K^+^ channels ^31,33,34^. Although dSPNs do express M_1_Rs, they only have been reported to modulate somatic K_v_7 channels ^33,34^, which alone had little effect on our simulations. What M_4_Rs do effectively is suppress D_1_ dopamine receptor signaling ^36^. Given that phasic activation of ChIs can induce dopamine release through terminal nAChRs ^44,45^, M_4_Rs might serve to limit the D1R-mediated enhancement of dSPN spiking in response to strong glutamatergic input – much like that produced by ChI activation of THIs/NGFIs.

From a network perspective, what do these simulations tell us? Tonically active ChIs are phasically accelerated by salient stimuli that promote set-shifting or transitions in movement syllables coded by ensembles of dSPNs and iSPNs ^5–11,46,47^. By engaging the THI/NGFI circuit and modulating postsynaptic ion channels, transient ChI bursting should create a temporal window in which the probability of spiking in response to cortical or thalamic glutamatergic inputs is increased in quiescent SPNs and suppressed in spiking SPNs. The suppression in ongoing spiking should be particularly prominent in active dSPNs. In so doing, it should promote transitions in striatal ensemble activity necessary for sequencing movement syllables. Given the asymmetry in the coordinated modulation of iSPNs and dSPNs, synchronous activity among ChIs could lead to preferential activation of iSPNs, serving to create a more prolonged pause in movement, an event that commonly accompanies presentation of a salient stimulus ^44^.

Finally, in PD models, ChI-coupling to GIs is attenuated. It has long been hypothesized that striatal cholinergic signaling is elevated in PD ^19^. One of the earliest treatments for PD were mAChR antagonists. Although several lines of study were consistent with this notion ^48^, recent work using genetically encoded optical sensors have provided a firmer foundation, showing a robust elevation in evoked striatal ACh release by ChIs in mouse models of late-stage PD ^20^. In agreement with the postulate that this increase largely reflected the loss of inhibitory, presynaptic D_2_R signaling in ChIs^20,49^, ACh release measured with the GRAB_ACh_ sensor also was elevated in premotor or prodromal MCI-Park mice where there was essentially complete loss of dopamine release in the DLS, but not parkinsonism. That said, these mice manifest deficits in complex motor behaviors and movement sequencing in the open field ^23^.

Although ChI release of ACh was elevated in models of both prodromal and late-stage PD, the ability of ChIs to drive the interneuron circuitry was profoundly impaired. Both fast THI-mediated and slow NGFI-mediated GABA_A_R currents were diminished in slices from PD models. This was true in both iSPNs and dSPNs. The deficit was attributable to down-regulation in the expression or surface positioning of nAChRs, as beta2 subunit mRNA abundance was reduced in both THIs and NGFIs from 6-OHDA-lesioned mice. Moreover, optogenetic activation of ChIs evoked significantly smaller currents in NGFIs following 6-OHDA lesioning. Interestingly, the reduction in nAChR-mediated currents was largely a consequence of a reduction in a slow component of the EPSC, which might reflect the engagement of extrasynaptic nAChRs. Lastly, directly activating NPY+ GABAergic interneurons, which include NGFIs and LTSIs, produced postsynaptic GABA_A_R-mediated currents that were indistinguishable in slices from control and 6-OHDA lesioned mice. Although THIs were not directly interrogated in these latter experiments, there is no obvious reason to doubt that they manifested changes similar to those in NGFIs. An unresolved question is whether the coupling of these GABAergic interneurons to cortical and thalamic circuits remains intact in the PD models.

Taken together, these results suggest that the dis-inhibition of ACh release early in the progression of PD triggers a compensatory down-regulation in nAChR mRNA level in THIs and NGFIs, resulting in an attenuation of ChI coupling to both iSPNs and dSPNs. As outlined above, this coupling deficit should impair the ability of ChIs to shift striatal ensemble activity in response to salient stimuli. Indeed, in prodromal, as well as parkinsonian MCI-Park mice, there is a significant disruption in the ability to perform sensory guided, sequential movements and in the ability to smoothly navigate a novel environment ^23^. In addition, deficits in set-shifting and behavioral flexibility have been noted in PD models and PD patients ^50–55^. While this constellation of disease symptoms could be due in part to alterations in cortical function, striatal determinants, including alterations in ChI regulation of striatal GABAergic circuitry, are also highly likely to play a role.

An obvious translational question is how this change in striatal function might be ameliorated or reversed. Although acute treatment of brain slices with dopaminergic agonists or levodopa administration to mice prior to sacrifice did not discernibly restore ChI coupling to SPNs in PD models (data not shown), chronic treatment may be more successful. This type of intervention could moderate ACh release and lead to a compensatory up-regulation in nAChR expression by THIs and NGFIs. Alternatively, it is possible that THIs and NGFIs interneurons express dopamine receptors that regulate nAChR expression and that restoring dopaminergic signaling will reverse the functional connectivity with ChIs. ACh release in PD models also could be moderated by restoration of autoreceptor function, which might be accomplished by suppression of RGS4 signaling in ChIs ^49^.

## Methods

### Animals

All animal experiments were conducted in accordance with the NIH Guide for the Care and Use of Laboratory Animals and were approved by the Northwestern University Institutional Animal Care and Use Committee (IACUC). The studies were supported under approved animal protocols IS00015064 (NIH/NINDS R37NS034696), IS00019822 (Aligning Science Across Parkinson’s [ASAP] grant ASAP-020551), and IS00010979 (Freedom Together Foundation grant GR-2021-2960). Northwestern University maintains an Animal Welfare Assurance on file with the Office of Laboratory Animal Welfare (OLAW; Assurance No. D16-00182/A3283-01), is registered with the United States Department of Agriculture (USDA Registration No. 33-R-0129), and is accredited by AAALAC International (File No. 000602).

For ex vivo experiments, C57BL/6 hemizygous mice expressing fluorescent reporters under the control of dopamine receptor regulatory elements were used. These included: *Drd1a*-tdTomato (RRID:MMRRC_030512-UNC); *Drd2*-eGFP (RRID:MMRRC_000230-UNC). Both lines were backcrossed to the C57BL/6 background. In some experiments, these mice were further crossed with: *ChAT*-IRES-Cre mice (B6;129S6-Chattm2(cre)Lowl/J, RRID:IMSR_JAX:006410) to selectively activate ChIs; *NPY*-Cre mice (RRID: IMSR_JAX:027851) to selectively activate NPY expressing GABAergic interneurons. To target NGFIs and record from SPNs, *Ndnf*-IRES-FlpO mice (B6(Cg)-Ndnftm1.1(flpo)Ispgl/J, RRID:IMSR_JAX:034876) were also used.

For Parkinson’s disease model experiments, MCI-Park mice were generated by crossing *Ndufs2*tm1.1Job Tg(CAG-cre/Esr1*)5Amc/J, (RRID:IMSR_JAX:038571) mice (on a mixed background: C57BL/6J, C57BL/6N, CBA/J, SWR, 129X1/SvJ, 129S1/Sv, and SJL) with DAT-IRES-cre (B6.SJL-*SlcCa3*tm1.1(cre)Bkmn/J, RRID:IMSR_JAX:006660) in which Cre expression is heterozygous.

To selectively activate ChI in MCI-Park mice, they were crossed with a *ChAT* Flpo mouse line that was generously provided by the Dauer laboratory (William T. Dauer; Peter O’Donnell Jr. Brain Institute, Department of Neurology, Department of Neuroscience, UT Southwestern), and Jay Li (Medical Scientist Training Program, Cellular and Molecular Biology Graduate Program, Department of Neurology, University of Michigan). This mouse line was generated by inserting an IRES-FLPo construct downstream of the Chat stop codon, similar to previously described Chat-Cre lines (RRID does not currently exist). Only male mice were used for experiments.

For CRISPR experiments, six-month-old ChAT-Cre × A2a-eGFP wt mice were used. These mice were generated by crossing ChAT-Cre mice (Jackson Laboratory; RRID:IMSR_JAX:006410) with A2a-eGFP mice (STOCK Tg(Adora2a-EGFP)EP141Gsat/Mmucd; RRID:MMRRC_010541-UCD), which had been previously backcrossed to C57BL/6J in the Surmeier laboratory.

### Stereotaxic brain surgery

Mice were anesthetized using a precision isoflurane vaporizer (SomnoSuite, Kent Scientific Corporation), a small-animal anesthesia system specifically designed for rodent procedures. Anesthesia was confirmed by the absence of reflexive responses to toe or tail pinch and the onset of complete immobility, accompanied by a stable respiratory rate of approximately one deep gasp per second, indicative of sufficient isoflurane-induced sedation for surgical manipulation. The animal was then secured in a stereotaxic frame (Model 940, David Kopf Instruments) equipped with a Cunningham adaptor (Harvard Apparatus) to ensure consistent anesthesia delivery throughout the procedure.

Following anesthetic induction, perioperative analgesia was provided. Pre-operatively, mice received a single subcutaneous loading dose of meloxicam (2 mg/kg; Meloxivet, Dechra, 5 mg/ml; NDC 17033-051-10), diluted per Northwestern University Center for Comparative Medicine (CCM) veterinary recommendations. The local anesthetic bupivacaine (2 mg/kg; Bupivacaine Hydrochloride Injection, 0.5%, 5 mg/ml, Eugia US; NDC 55150-0250-50) was infiltrated at the incision site. Post-operatively, mice received a single dose of extended-release buprenorphine (1 mg/kg; Buprenorphine ER, 0.5 mg/ml, Wedgewood Connect; NDC 79926-057-17). When buprenorphine ER was unavailable, meloxicam was continued at a maintenance dose of 1 mg/kg every 24 h for 1–2 days. The scalp was then sterilized, the fur removed, and an incision made to expose the skull. A small craniotomy was drilled at the targeted stereotaxic coordinates for the DLS: anterior-posterior (AP) +0.7 mm relative to bregma, medial-lateral (ML) −2.3 mm, and dorsal-ventral (DV) −3.30 mm from the dura. These coordinates were determined using the Allen Mouse Brain Atlas (online version 1, 2008; RRID:SCR_002978; http://mouse.brain-map.org/static/atlas).

For viral injections, approximately 600 nL of adeno-associated virus (AAV) was loaded into a pulled glass micropipette (Drummond Scientific; pulled using a P-97 puller, Sutter Instruments) and delivered unilaterally into the DLS. Following injection, the needle was left in place for 5 minutes to allow for adequate tissue absorption of the viral solution before being slowly withdrawn to minimize backflow and ensure targeted delivery (https://dx.doi.org/10.17504/protocols.io.81wgby191vpk/v1). Except where otherwise noted, Cre-dependent optogenetic activation used a Chronos opsin (AAV-EF1α-FLEX-rc [Chronos-GFP]; RRID:Addgene_62725).

### 6-OHDA lesioning

Unilateral lesions of the nigrostriatal dopaminergic (DA) pathway were induced by stereotaxic injection of 6-hydroxydopamine (6-OHDA) into the MFB. The neurotoxin 6- OHDA was freshly prepared at a concentration of 2.5 µg/µl in 0.04% ascorbic acid (to prevent oxidation) and administered at a total dose of 2.5 µg in a volume of 1 µl. Injections were delivered using calibrated glass micropipettes. Stereotaxic coordinates used to target the MFB were: (AP) −0.7 mm from bregma, (ML) +1.2 mm, and (DV) −4.75 mm from the dura. The toxin was injected at a controlled rate of 0.1 µl/min by manually applying gentle positive pressure to the pipette using a handheld syringe assembly. Then, the pipette was left in place for an additional 15-20 minute post-injection to allow sufficient diffusion and tissue retention of the solution, and to minimize reflux along the injection tract. To assess the extent of nigrostriatal dopaminergic (DA) neuron damage, a limb-use asymmetry test (cylinder test) was regularly performed before each electrophysiological experiment. Mice were placed individually into a transparent glass cylinder (8 cm in diameter, 12 cm in height), and spontaneous forelimb use was recorded for a duration of 5 minutes during exploratory rearing and wall contacts. During this time, the number of independent wall touches made by the unimpaired (ipsilateral) and impaired (contralateral) forelimbs, as well as simultaneous use of both limbs, was quantified. The percentage of impaired (contralateral) forelimb use was calculated relative to the total number of forelimb movements. Mice with successful 6-OHDA lesions typically showed impaired limb usage ranging from 5% to 30%, indicating effective unilateral disruption of the nigrostriatal pathway. (https://dx.doi.org/10.17504/protocols.io.kqdg3q5w1v25/v1).

### Stereotaxic injection of AAVs

A computer-assisted stereotaxic system (Leica Biosystems, Buffalo Grove, IL) was used to deliver viral injections into the striatum. Mice were anesthetized with isoflurane, and two striatal injection sites were targeted using the following coordinates (relative to bregma): Site 1: lateral, 2.15 mm; posterior, 0.98 mm; depth, 2.8 mm; Site 2: lateral, 2.0 mm; posterior, 0.74 mm; depth, 2.6 mm.

For GABA_A_R delta subunit knockdown experiments and corresponding controls, viruses were co-injected with the Chronos opsin. At each site, 500 nL of virus was injected using an IM 300 microinjector (Narishige, Japan) at a moderate injection pressure (6–8 psi) and duration (60–300 s). Electrophysiological recordings were performed five weeks post-injection (https://dx.doi.org/10.17504/protocols.io.81wgby191vpk/v1).

### GABA_A_R knockdown experiments

CRISPR technology was used to investigate the involvement of the GABA_A_R receptor delta subunit (GABAδ) in disynaptic GABAergic IPSCs following ChI optogenetic activation. Six-month-old ChAT-Cre × A2a-eGFP wt mice with a C57BL/6 background were used in these experiments. These mice were generated by crossing ChAT-Cre mice (Jackson Laboratory; RRID:IMSR_JAX:006410) with A2a-eGFP mice (RRID:MMRRC_010541-UCD), which had been previously backcrossed to C57Bl/6J in the Surmeier laboratory.

To knockdown GABAδ, a mixture of the guide RNA virus (AAV9-Syn-FRed-U6gRsGABAd, 2E+13 vg/ml, Virovek; RRID:Addgene_258848) and the SaCas9 virus (AAV9-Syn-SaCas9s-mWPRE-hGHPolyA / pFB-Syn-SaCas9s-mWPRE-hGHpA, 2E+13 vg/ml, Virovek; RRID:Addgene_245385) were co-injected in the dorsal striatum. In the control experiments, only the guide RNA virus was injected. For the optogenetic activation of the ChIs, the dorsal striatum of ChAT-Cre X A2a-eGFP wt mice was injected with a Chronos opsin (AAV9-Syn-FLEX-Chronos-GFP, 2E+13 vg/ml, Virovek; RRID:Addgene_62722).

For optogenetic activation of NGF/NDNF interneurons, the dorsal striatum of *Ndnf*-Flp (RRID:IMSR_JAX:034876) × A2a-eGFP mice (RRID:MMRRC_010541-UCD) was injected with a Flp-dependent opsin virus (AAV8-CAG-FLPX-rc [ChrimsonR-tdTomato]; RRID:Addgene_130909). The same Flp-dependent Chrimson virus (RRID:Addgene_130909) was used for ChI activation in prodromal MCI-Park mice.

To assess the functional connectivity of ChIs with NGFIs, *ChAT*-Cre mice (RRID:IMSR_JAX:006410) were crossed with *Ndnf*-IRES-FlpO mice (RRID:IMSR_JAX:034876). The DLS of these mice was co-injected with a Cre-dependent Chronos opsin (AAV-EF1α-FLEX-rc [Chronos-GFP]; RRID:Addgene_62725) and a Flp-dependent reporter (AAV-Ef1a-fDIO-mCherry; RRID:Addgene_114471).

### Brain slice preparation

Mice were deeply anesthetized via intraperitoneal injection of a ketamine (100 mg/kg) and xylazine (7 mg/kg) mixture. Once fully anesthetized, animals were transcardially perfused with ice-cold, oxygenated cutting solution composed of (in mM): 125 NaCl, 2.5 KCl, 1.25 NaH_2_PO_4_, 0.5 CaCl_2_, 2.0 MgCl_2_, 26 NaHCO_3_, and 10.1 glucose (305 mOsm/L, pH 7.3). Coronal brain slices (300 μm thick) containing Striatum were prepared using a vibratome (VT1200S, Leica Biosystems, Wetzlar, Germany).

Following slicing, sections were incubated at 34 °C for 40 minutes in artificial cerebrospinal fluid (ACSF) consisting of (in mM): 125 NaCl, 2.5 KCl, 1.25 NaH_2_PO_4_, 2.0 CaCl_2_, 1.0 MgCl_2_, 26 NaHCO_3_, and 10.1 glucose. Slices were then maintained at room temperature for at least 30 minutes before electrophysiological recording to allow for recovery and acclimation. Throughout the entire procedure, all solutions were continuously bubbled with carbogen gas (95% O_2_ / 5% CO_2_) to ensure optimal oxygenation and pH balance (https://dx.doi.org/10.17504/protocols.io.dm6gp328jvzp/v2).

### Slice Electrophysiology

Individual brain slices were transferred to a recording chamber and continuously superfused with oxygenated ACSF at a rate of 2–3 ml/min at room temperature. D_1_-tdTomato– or D_2_-eGFP–expressing SPNs in the striatum were visually identified using an upright Olympus BX-51WI microscope equipped with a 40X/0.8 NA water-dipping objective lens and a CCD video camera (USB 3.0 Digital Camera, Thorlabs). Whole-cell or perforated patch-clamp recordings were performed from fluorescently labeled SPNs and NGFIs. Patch pipettes (3–4 MΩ resistance) were filled with an internal solution containing (in mM): 140 CsMeSO_3_, 3 NaCl, 0.25 EGTA, 10 HEPES, 2 Mg-ATP, 0.3 Na-GTP, 10 TEA-Cl, and 5 QX-314 (pH adjusted with CsOH to 7.3, 290–300 mOsm/L). Cells were voltage-clamped at −80 mV. For cell-attached recordings of ChIs, cells were identified by both their fluorescence and the presence of spontaneous firing activity. Recordings were performed using a MultiClamp 700B amplifier (Axon Instruments), filtered at 2 kHz, and digitized at a sampling rate of 10 kHz using Clampex 10.7 software (RRID:SCR_011323). Series resistance was below 30 MΩ and was continuously monitored and the holding current did not exceed approximately −300 pA. PSCs were evoked once per minute using a single 5-ms LED light pulse (X-Cite, Excelitas), illuminating the whole field. To isolate GABAergic PSCs, ionotropic and metabotropic glutamatergic and GABA_B_ receptor antagonists were used (10 µM NBQX, 10 µM D-AP5, 50 µM CPCCOEt, 1 µM MPEP hydrochloride, 2 µM CGP55845) (https://dx.doi.org/10.17504/protocols.io.rm7vzx1w2gx1/v2).

### Two-Photon Imaging of ACh Release

ACh dynamics were evaluated using the genetically encoded ACh sensor GRAB_ACh3.0_, delivered as AAV9-hSyn-GRAB_ACh3.0 / pAAV-hSyn-GRAB_ACh3.0 (RRID:Addgene_121922), expressed in the DLS and imaged via two-photon laser scanning microscopy (2PLSM). Acute coronal brain slices containing striatal GRABACh3.0 expression were prepared as previously described, then transferred to a submerged recording chamber and continuously superfused with oxygenated ACSF maintained at room temperature. Imaging was performed using a Chameleon Ultra II two-photon laser (Coherent, Santa Clara, CA), tuned to an excitation wavelength of 920 nm. Fluorescence emission from GRABACh3.0 was captured using an Ultima In Vitro Multiphoton Microscope system (Bruker, Billerica, MA) equipped with a 60x/1.0 NA water-immersion objective (Olympus) and a GaAsP photomultiplier tube (Hamamatsu H7422P-40; 490–560 nm detection range). Time-series images were acquired with a pixel resolution of 0.388 μm × 0.388 μm, 4-μs dwell time per pixel, and a frame rate of 21.26 frames per second using Galvo spiral scanning mode (Prairie View 5.3, Bruker; RRID:SCR_017142). After a 5-second baseline acquisition period, synchronous ACh release was evoked by electrical stimulation using a concentric bipolar electrode (CBAPD75, FHC) placed approximately 150 μm dorsal to the imaging field. Stimulation protocol included a single 1-ms pulse at 0.3 mA. Image acquisition continued for at least 5 seconds following stimulation. Three stimulation trials were recorded for each region of interest (ROI) and data was mean averaged. Fluorescence intensity (F) was first background-subtracted (the background noise resulted from PMT, determined under zero laser power at same PMT voltage), and the baseline fluorescence (F₀) was defined as the mean intensity during the 1 second preceding stimulation. The ΔF (F − F₀) or the amplitude of the response, was normalized to the baseline fluorescence (F₀) using the formula: ΔF_norm = (F − F₀) / (F₀). An unpaired t-test was then performed to compare the distribution of data obtained from MCI-Park mice and their respective controls (https://dx.doi.org/10.17504/protocols.io.n92ldry68g5b/v1).

### RNAscope In Situ Hybridization

ChAT-Cre mice were sacrificed four weeks after unilateral 6-OHDA lesioning. Brain tissues were rapidly collected, embedded in Optimal Cutting Temperature (OCT) compound, and flash frozen. Coronal brain sections (12 μm) were prepared using a cryostat, and slides were stored at −80 °C until use. RNAscope in situ hybridization assays were performed on fresh frozen sections using the RNAscope Multiplex Fluorescent Detection Kit v2 (ACD/Bio-Techne, Cat#323110), according to the manufacturer’s protocol. The following RNAscope probes were used: Mm-CHRNb2 (ACD/Bio-Techne, Cat#449231) targeting nicotinic acetylcholine receptor beta 2 subunit (CHRNB2); Mm-NDNF-C3 (ACD/Bio-Techne, Cat. #447471-C3) targeting neuron-derived neurotrophic factor (NDNF); and Mm-TH-C2 (ACD/Bio-Techne, Cat#317621-C2) targeting tyrosine hydroxylase (TH).

Slides were mounted with ProLong™ Diamond Antifade Mountant containing DAPI (Invitrogen) and imaged using an Olympus FluoView FV10i confocal laser scanning microscope. To quantify CHRNB2 expression in NDNF- or TH-expressing neurons, cells in the dorsal striatum were outlined in FluoView FV10-ASW (Olympus, RRID:SCR_014215) to define regions of interest (ROIs), and CHRNB2-positive signal particles within each ROI were counted manually. Data are presented as CHRNB2 signal particles per cell and compared between the control and lesion sides. For each condition, 39–45 cells from four mice were analyzed (https://dx.doi.org/10.17504/protocols.io.14egn3odml5d/v1).

### Immunocytochemistry

Mice were deeply anesthetized and transcardially perfused with 0.9% saline, followed by 4% paraformaldehyde prepared in 0.1 M phosphate buffer. Brains were removed and incubated overnight at 4 °C in 4% paraformaldehyde. Serial coronal sections (80 μm thickness) encompassing the striatum were collected. For immunofluorescence labeling, free-floating sections were incubated with a mouse monoclonal antibody against tyrosine hydroxylase (TH; ImmunoStar, Cat#22941; RRID:AB_572268) diluted 1:1,000 in blocking solution and maintained at 4 °C for 24 h. Sections were then washed in phosphate-buffered saline (PBS) and incubated with a goat anti-mouse Alexa Fluor 594–conjugated secondary antibody (Thermo Fisher Scientific/Invitrogen, Cat#A-11005; RRID:AB_2534073; 1:500) for 30 min at room temperature. Following final washes, sections were mounted using ProLong™ Diamond Antifade Mountant containing DAPI (Thermo Fisher Scientific, Cat#P36962). Fluorescence images were acquired using a laser-scanning confocal microscope (Olympus FV10i) equipped with ×10/0.4 NA (dry) and ×60/1.35 NA (oil immersion) objectives (https://dx.doi.org/10.17504/protocols.io.bp2l6xpr1lqe/v1). Fiji (NIH, RRID:SCR_002285) was used to adjust images for brightness, contrast, and pseudo-coloring.

### Independent study (Fig. 5S)

Experiments shown in Supplementary Figure S5 were performed independently. Adult ChAT-ChR2-eYFP mice (B6.Cg-Tg(Chat-COP4*H134R/EYFP, Slc18a3)6Gfng/J; Jackson Laboratory; RRID:IMSR_JAX:014546) were used for all experiments. Unilateral dopaminergic lesions were induced by stereotaxic injection of 6-hydroxydopamine (6-OHDA; 3.5 mg/mL in 0.1% ascorbic acid) into the substantia nigra pars compacta (SNc; AP −2.7 mm, ML +1.35 mm, DV −4.2 mm from bregma). A total volume of 1.3 μL was delivered using a Nanoject II Auto-Nanoliter Injector. The contralateral hemisphere received saline and served as an internal control.

Mice were anesthetized with ketamine (80 mg/kg) and xylazine (20 mg/kg) and transcardially perfused with ice-cold, oxygenated NMDG-based cutting solution. Coronal brain slices (300 μm thick) containing the striatum were prepared using a VT1200S vibratome (Leica Microsystems). Following slicing, sections were incubated in oxygenated NMDG solution at 35°C for 5 minutes and then transferred to oxygenated ACSF at room temperature until recording. During recordings, slices were continuously perfused with oxygenated ACSF maintained at 32–34°C.

Whole-cell voltage-clamp recordings were obtained from visually identified striatal SPNs using infrared differential interference contrast microscopy. Recording electrodes (3–5 MΩ) were filled with a CsCl-based internal solution containing biocytin for post hoc morphological verification. Signals were amplified using a MultiClamp 700B amplifier and digitized with an ITC-1600 interface. Recordings were acquired using Axograph X software (RRID:SCR_014284). ChR2-expressing ChIs were activated using 2-ms pulses of blue light (450 nm) delivered every 30 s.

### Modeling

Simulations used a NEURON (Neuron 8.2.6; RRID:SCR_005393) with Python (Python Programming Language RRID:SCR_008394) model of morphologically reconstructed striatal SPNs, implemented as an updated version of Day et al. ^16^ and integrated into previously established SPN modelling frameworks ^29,56,57^. Cytoplasmic resistivity (Ra) was set to 200 Ωcm and specific membrane capacitance was 1 µFcm⁻^2^. The models comprised biophysically detailed representations of a dSPN and iSPN consisting of 822 and 710 segments, respectively, incorporating the following active and passive conductances: transient fast inactivating Na^+^ (Naf), persistent Na^+^ (Nap), fast A-type K^+^ (K_v_4.2), slowly inactivating K^+^ (K_v_1.2), inwardly-rectifying K^+^ (K_ir_), delayed rectifier K^+^ (K_v_3.1/3.2), KCNQ (K_v_7), small conductance Ca^2+^-activated K^+^ (SK), large conductance Ca^2+^-activated K^+^ (BK), L-type Ca^2+^ (Ca_v_ 1.2 and 1.3), N-type Ca^2+^ (Ca_v_ 2.2), R-type Ca^2+^ (Ca_v_ 2.3), T-type Ca^2+^ (Ca_v_3.2 and 3.3). Channel distributions across cellular compartments were as previously described based on Lindroos et al. ^56^.

Synaptic spines were added to all dendritic locations further than 30 µm from the soma at a density of 1.61 per µm, yielding approximately 5300 and 4400 spines for the reconstructed dSPN and iSPN, respectively. Spines consisted of a cylindrical head with a diameter of 0.5 µm connected to the dendrite via a neck 1 µm in length with a diameter of 0.1 µm. The reconstructed dSPN model had a resting membrane potential of −85 mV. A hyperpolarizing current step of 200 pA produced a rectified range input resistance of approximately 85 MΩ and a membrane time constant of 10.5 ms. The estimated whole cell capacitance was approximately 180 pF for the dSPN and 160 pF for the iSPN model. Candidate spines were selected as separate nearest neighbors along a dendrite, beginning approximately two thirds of the distance from the soma. Glutamatergic synaptic input was delivered via α-amino-3-hydroxy-5-methyl-4-isoxazolepropionic acid (AMPA) and N-methyl-D-aspartate (NMDA) conductances inserted into these spines. Synaptic currents were modeled using a two-state kinetic scheme in which the normalized peak conductance was determined by two time constants T_1_ and T_2_ using an equation of the form (1 − e^−t/τ1^) e^−t/τ2^, where T_2_ is greater than T_1_, as previously described ^29,56,57^. The maximal conductances of AMPA and NMDA responses were 350 and 752.5 pS, respectively, with reversal potentials set to 0 mV. Spines were activated sequentially at 1 ms intervals with stimulation progressing toward the soma. Subthreshold and suprathreshold conditions were determined by the number and spatial distribution of activated synapses. Spines were activated as clustered nearest neighbors along a dendritic segment, with activation proceeding sequentially at 1 ms intervals in the distal to proximal direction within each site. Where multiple dendritic sites were stimulated, clustered inputs at each site were activated simultaneously. GABAergic conductances, including synaptic and extrasynaptic receptors, were inserted onto dendritic locations and activated simultaneously. The maximal conductance was 1000 pS and the reversal potential was set to −60 mV. Fast synaptic and slow extrasynaptic GABAergic activation regimes were implemented by varying receptor number and kinetics. In simulations where clustered glutamatergic input alone was subthreshold, overall conductance was varied to produce the same local dendritic postsynaptic potential at the site of clustered glutamatergic input. In simulations where clustered glutamatergic input alone was suprathreshold, overall conductance loads were fixed within each condition, such that fast synaptic and slow extrasynaptic GABAergic activation produced different postsynaptic potential amplitudes due to their distinct kinetics. Specific conductance loads and receptor configurations are detailed in the corresponding figures. Temporal interactions between excitation and inhibition were examined by pairing glutamatergic and GABAergic inputs at defined temporal offsets (Δt = t_GLUT_ − t_GABA_), where negative values indicate that GABAergic activation preceded glutamatergic input. The range and resolution of Δt values varied across simulation protocols and are indicated in the corresponding figures. For iSPNs, M_1_ muscarinic receptor activation was simulated by modulation of K_ir_2, K_v_4 and K_v_7 K^+^ channels. For dSPNs, this modulation was limited to K_v_7 K^+^ channels^31,33,34^. Modulation was modelled as 50% reduction in each conductance for channels affected. Simulations were performed under paired control and M_1_ modulated conditions to assess the effects of cholinergic signaling on dendritic integration and neuronal output. Simulation output measures included dendritic postsynaptic potential amplitude, supralinear integration relative to glutamatergic input alone, dendritic spike generation, and somatic spike output depending on the stimulation regime. All code for the simulations is publicly available (https://github.com/vernonclarke/msNEURON_Belal2026; https://doi.org/10.5281/zenodo.20705696).

### Statistics

PSC waveforms were fit by nonlinear least squares in R (versions 4.4 - 4.6; R Project for Statistical Computing; RRID:SCR_001905) using product functions of the form A(1 − e^−t/τ1^) e^−t/τ2^. Amplitudes were parameterized as peak amplitudes, with rise and decay time constants derived from the fitted parameters (T_rise_ = T_1_ T_2/_(T_1_ + T_2_); T_decay_ = T_2_). Two-component fits were used throughout to decompose PSCs into fast and slow components with corresponding amplitudes, charge transfer, and decay time constants. In a subset of analyses, one- and two-component fits were compared per recording using the Bayesian information criterion (BIC) to verify whether a two-component model provided a better description of that experimental data. Two-component fitted values were occasionally screened for bivariate outliers in fast/slow PSC amplitude (A_fast_, A_slow_) using robust Mahalanobis distances (MCD estimator) against a conservative χ^2^ threshold that resulted in a Mahalanobis cutoff ~ 5.4, following the method of Rousseeuw and van Driessen, 1999 ^58^.

For electrophysiological measurements, including PSC amplitude, charge transfer, and decay time constants, normality was not assumed and hypothesis testing used nonparametric Wilcoxon tests. Paired within-recording comparisons, including baseline versus drug condition and fast versus slow components from the same recording, were analyzed using Wilcoxon signed-rank tests. Independent between-group comparisons, including dSPN versus iSPN and control versus 6-OHDA groups, were analyzed using Wilcoxon rank-sum tests. Tests were two-sided with α = 0.05. Exact *p* values were used when available; otherwise the normal approximation was applied. Where multiple comparisons were made within a logical family of tests, *p* values were adjusted using the Holm’s step-down method. A family was defined as all comparisons of a given measure within a particular experimental dataset. So, for example, amplitude comparisons in a given figure panel were corrected together, separately from charge-transfer or decay-constant comparisons. Adjusted *p* values are reported throughout; the unadjusted values are provided in the source data.

For fluorescence imaging data (GRAB_ACh3.0_ sensor ΔF/F₀, Figure 7) and RNAscope puncta counts (Figure 11), observations were nested within slices, fields, and/or animals, and mixed-effects models were used. For Figure 7, the outcome was modelled as a function of condition with a random intercept for slice. For Figure 11, RNAscope counts were modelled as a function of group with random intercepts for field nested within animal. Robust mixed-effects models were fit (robustlmm), and fixed effects were evaluated by cluster bootstrap. For Figure 7, slices were resampled with replacement (with 9,999 bootstraps); for Figure 11, animals were resampled with replacement while preserving nested fields (with 1,999 bootstraps). Bootstrap *p* values were calculated from the empirical distribution of fixed-effect estimates. Figures and statistical summaries were generated using custom R scripts (https://github.com/vernonclarke/analysis_Belal2026; https://doi.org/10.5281/zenodo.20658501).

## Acknowledgements

The authors thank Chrissy Weber-Schmidt, Sasha Ulrich and David Wokosin for their technical assistance and support. This research was funded by grants to DJS from Aligning Science Across Parkinson’s [ASAP020551] through the Michael J. Fox Foundation for Parkinson’s Research (MJFF); Aligning Science Across Parkinson’s Collaborative Research Network, Chevy Chase, MD, 20815, https://parkinsonsroadmap.org; the Freedom Together Foundation [GR-2021-2960], 875 Third Avenue, 29th Floor, New York, NY 10022, https://www.freedomtogether.org/ and the National Institute of Neurological Disorders and Stroke [R37 NS034696], P.O. Box 5801. Bethesda, MD 20824; https://www.ninds.nih.gov. The funders had no role in study design, data collection and analysis, decision to publish, or preparation of the manuscript. For the purpose of open access, the authors have applied a CC BY public copyright license to all Author Accepted Manuscripts arising from this submission.

## Data availability

The data, protocols, and key lab materials generated in this study are listed in the Key Resource Table available at https://doi.org/10.5281/zenodo.20617251 alongside their persistent identifiers. Protocols are available at the protocols.io DOIs cited in the Methods. Raw and primary-processed neurophysiology, imaging, and RNAscope data are available through DANDI (DANDI:001832) located at https://doi.org/10.48324/dandi.001832/0.260611.2102. These data are formatted as Neurodata Without Borders (NWB; RRID:SCR_015242) files organized according to the Brain Imaging Data Structure (BIDS; RRID:SCR_016124) standard and released under a CC BY 4.0 license. Cleaned and summary data are available at https://doi.org/10.5281/zenodo.20029446 and are likewise released under a CC BY 4.0 license.

## Code availability

All custom code generated for this study is publicly available. Analysis code, providing a complete workflow from the raw DANDI datasets through processed data, statistical analyses, nonlinear least-squares fits, and final graphical outputs used in the manuscript, is available at https://github.com/vernonclarke/analysis_Belal2026 with an archived release at https://doi.org/10.5281/zenodo.20658501. Simulation code, providing a complete workflow to generate, analyze, and produce the final graphical outputs for all simulation figures in the manuscript, is available at https://github.com/vernonclarke/msNEURON_Belal2026 with an archived release at https://doi.org/10.5281/zenodo.20705697. The code is released under the MIT license.

## Author contributions

Conceptualization: M.B., V.R.J.C. and D.J.S.

Methodology: M.B., T.T., J.L., W.D., V.R.J.C. and D.J.S.

Software: V.R.J.C. Validation: E.B.G. and S.K. Formal analysis: V.R.J.C., Z.X.

Investigation: M.B., T.P.-R., E.B.G., S.K., Z.X. and E.I.

Resources: T.T., J.L., W.D. and D.J.S.

Visualization: V.R.J.C.

Writing: M.B., V.R.J.C. and D.J.S.

Editing: all authors.

Supervision: W.D., M.A., J.M.T. and D.J.S.

Funding acquisition: D.J.S.

---

**Figure S1.**
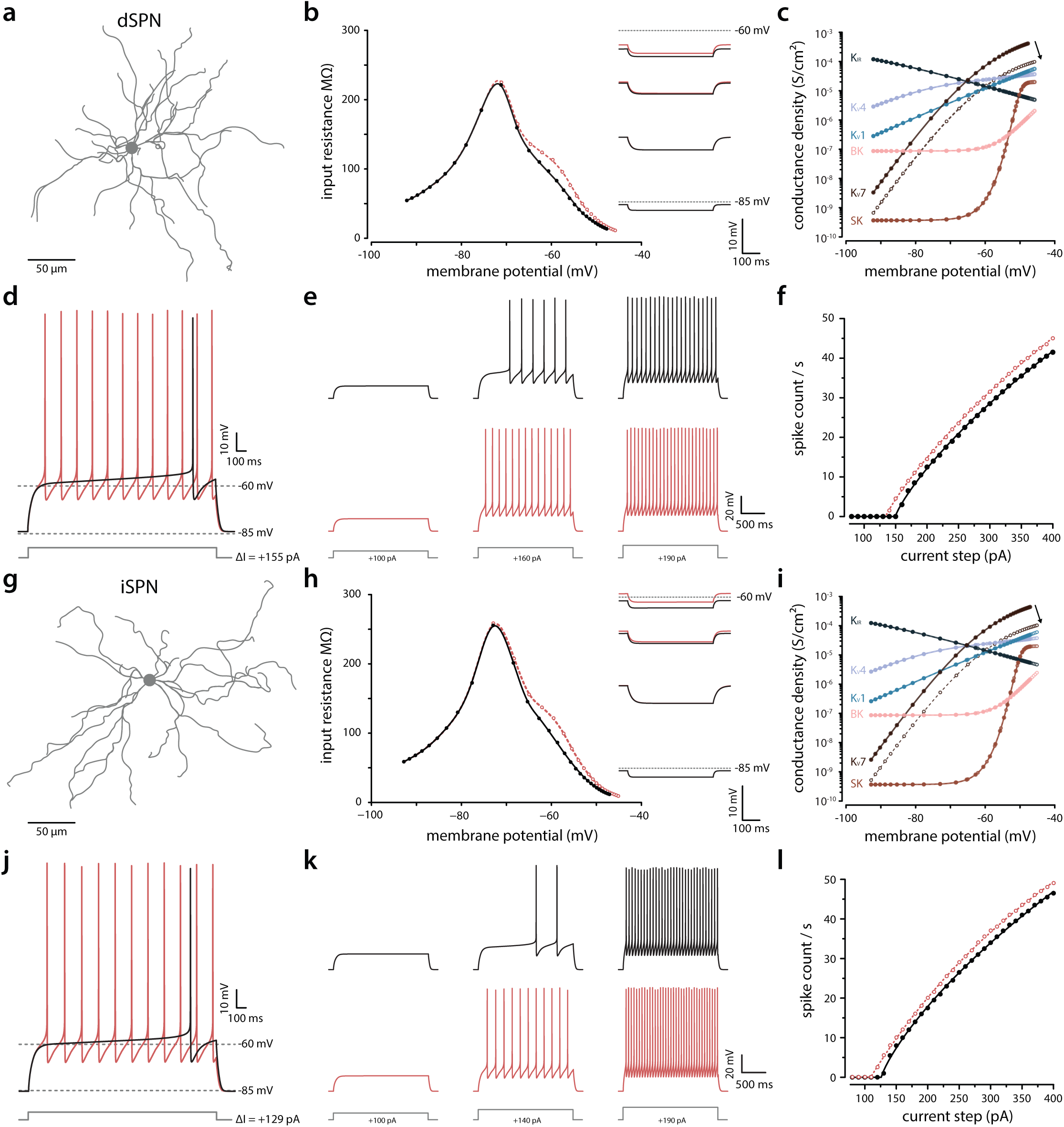
(a) Morphology of a reconstructed dSPN. (b) Plot of input resistance (measured using hyperpolarizing current steps of −10 pA, 1000 ms) as a function of holding membrane potential under control conditions (black, filled circles) and following partial block of K_v_7 potassium channels (80% reduction in local membrane conductance density where expressed; red, open circles). (c) Plot of membrane conductance density versus holding membrane potential. At hyperpolarized potentials, input resistance is dominated by high K_ir_2 conductance density. The increase in input resistance with depolarization reflects the intrinsic inward rectification of this channel, manifested as a reduction in K_ir_2 conductance density. At more depolarized potentials, the subsequent decrease in input resistance is primarily attributable to activation of other voltage-gated K⁺ channels, notably K_v_4 and Kv1 increasing their conductance densities. K_v_7 blockade only shifts its conductance density (open circles; direction indicated by an arrow). (d) Rheobase (defined as the minimum positive current to cause an action potential from membrane resting potential) was +155 pA under control (black line). The same current step causes a dramatic increase in spiking to the same current step following partial block of K_v_7 potassium channel (80% reduction; red line). (e) Single examples of increasing positive current steps and (f) summary to show the reduced rheobase and increased spike count rate to depolarizing current steps when K_v_7 is blocked (control: black, filled circles; K_v_7 blockade: red, open circles). Similar plots are shown for the simulated iSPN (g-l). The effects are qualitatively similar. The main differences are that input resistance is slightly elevated manifesting as an increased excitability / reduced rheobase (iSPN vs dSPN: +129 pA vs +155 pA).

**Figure S2.**
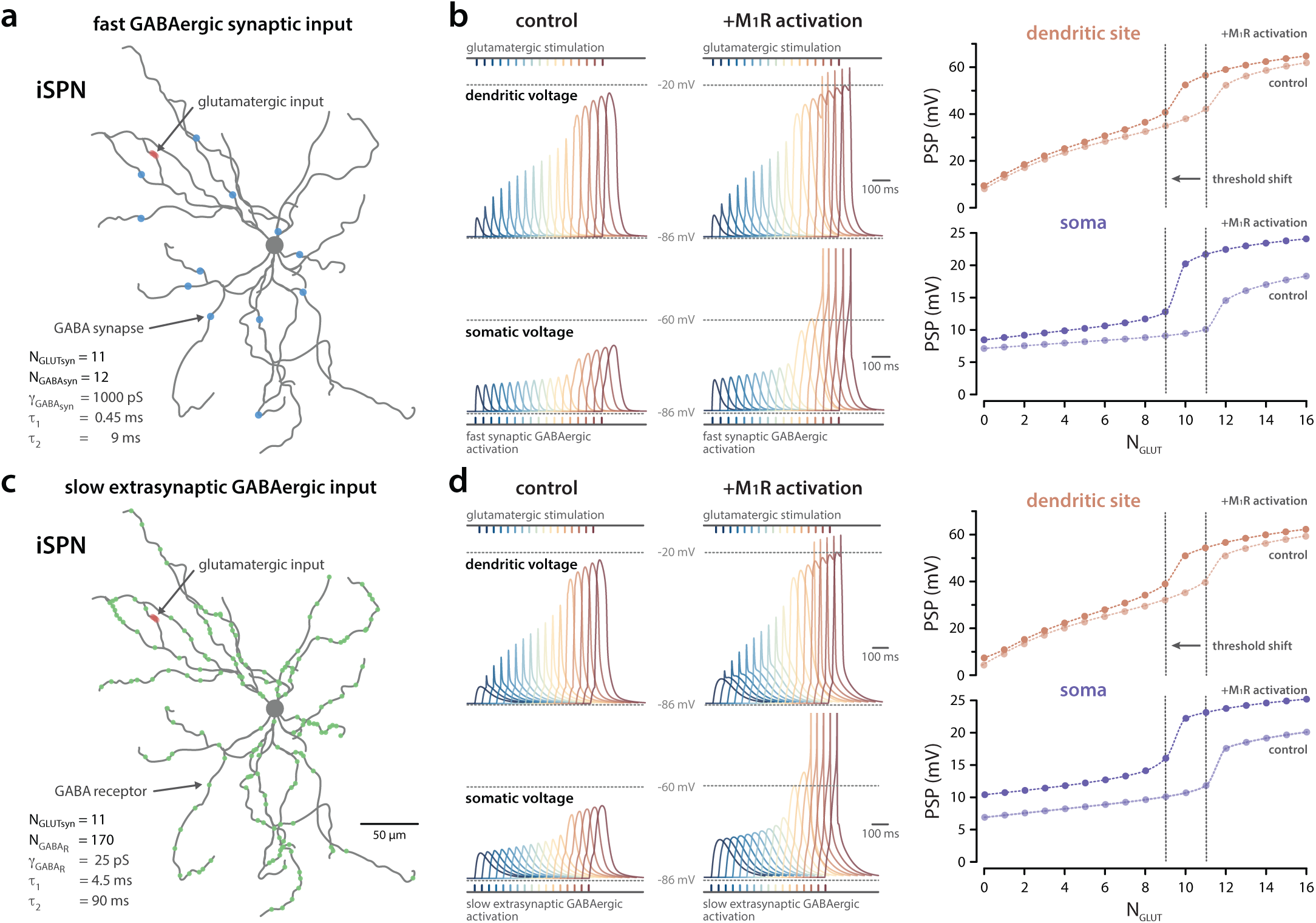
Effect of M_1_ mAChR activation on the temporal interaction between subthreshold glutamatergic and GABAergic inputs in a model iSPN. (a) Reconstructed iSPN morphology showing clustered glutamatergic input (red) and fast synaptic GABAergic input (blue). (b) Effect of increasing glutamatergic synapse number at a fixed dendritic site with fast synaptic GABAergic input (12 synapses each comprising 40 receptors with 25 pS peak conductance; Σg_GABA_ = 12 nS; PSP amplitude 8.0 mV at dendritic spike site) in control (left) and M_1_ receptor modulation of K_ir_2, K_v_4 and K_v_7 K^+^ channels (right). Color traces represent incremental glutamatergic recruitment (N_GLUT_ = 0-16) with fixed GABAergic activity (Δt = +10 ms; *i.e*. glutamate activity follows GABAergic input) at dendritic site of spike generation and cell soma. **M_1_** receptor activation lowers dendritic spike threshold from N_GLUT_ = 11 to 9. (c) Reconstructed iSPN morphology showing dendritic sites of clustered glutamatergic activation (red) and slow (green) extrasynaptic GABAergic receptor inputs. (d) Illustrating the effect of the slow GABAergic activation (170 receptors, 25 pS; Σg_GABA_ = 4.25 nS; PSP = 7.3 mV at dendritic spike site) on the same dendritic glutamatergic inputs as before (i.e. panel B; Δt = +30 ms). Again, M_1_ receptor activation lowers dendritic spike threshold from N_GLUT_ = 11 to 9.

**Figure S3.**
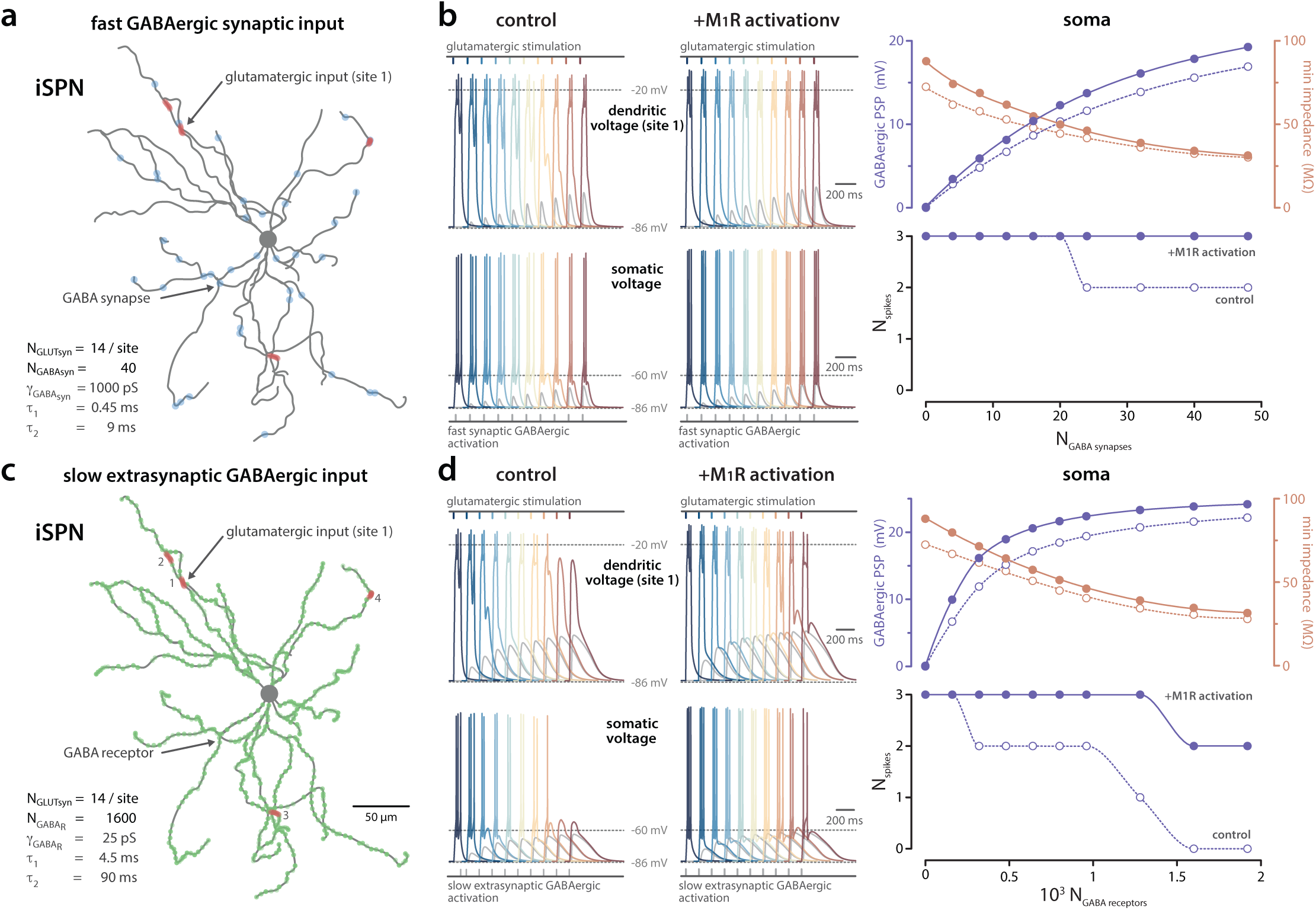
Effect of M_1_ mAChR activation on the interaction between suprathreshold glutamatergic input and varying amplitude, temporally fixed GABAergic input in a model iSPN. (a) Reconstructed iSPN morphology showing four dendritic sites of clustered glutamatergic activation (red; sites 1-4) and fast synaptic GABAergic input (blue). (b) Synaptic potentials at a dendritic site (site 1) and soma showing the effect of varying fast synaptic GABAergic input alone (gray) and when delivered 30 ms after suprathreshold glutamatergic input (14 inputs/site; color traces). Traces illustrate the effect of increasing GABAergic synaptic activity (0 - 48 synapses; each comprising 40 receptors with 25 pS peak conductance; blue to red) delivered at the same temporal offset (*i.e.* Δt = −30 ms) in the absence and presence of M_1_R activation. GABAergic synapse recruitment increased somatic depolarization (blue open circles; dotted line) with a concomitant decrease in impedance (orange open circles; dotted line). A small reduction in the total number of spikes generated at the soma was observed (blue open circles; dotted line). M_1_ receptor modulation of K_ir_2, K_v_4 and K_v_7 K^+^ channels increased GABAergic-mediated somatic depolarization and maintained somatic spike output (closed circles; solid line). (c, d) As in (a, b) but for slow extrasynaptic GABAergic receptor activation (0 - 1920 receptors; blue to red) color-matched to the same overall conductance loads as panel c (*i.e.* Σg_GABA_) at the same temporal offset *(i.e.* Δt = −30 ms). Extrasynaptic GABAergic receptor recruitment increased somatic depolarization (blue open circles; dotted line) with a concomitant decrease in impedance (orange open circles; dotted line). The suppression of spiking and its relief by M_1_R activation were more pronounced with slow GABAergic activation than fast synaptic activation at the same overall conductance load.

**Figure S4.**
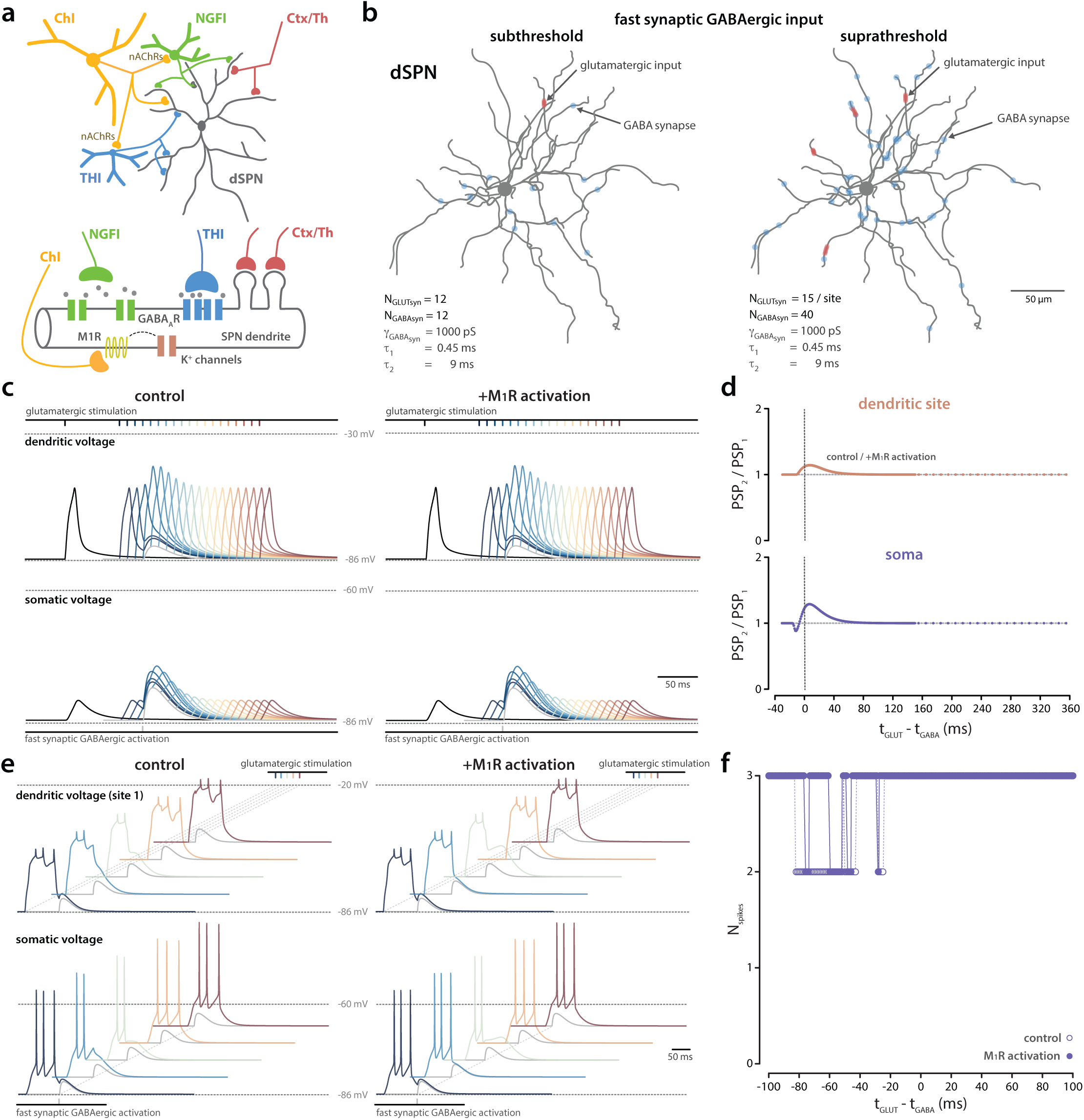
Effect of M_1_ mAChR activation on the interaction between glutamatergic input and fast synaptic GABAergic responses in a model dSPN. (a) Circuit diagram depicting cholinergic control of GABAergic microcircuits targeting SPNs and a schematic illustrating inputs onto a stretch of SPN dendrite. (b) Reconstructed dSPN morphology showing subthreshold (1 site) and suprathreshold (4 sites) clustered glutamatergic activation (red) and fast synaptic GABAergic input (blue). **(c)** Effect of fast synaptic GABAergic activation on glutamatergic synaptic potentials at dendritic and somatic sites in control (left) and following M_1_ receptor modulation of K_v_7 channels (right). Black trace: subthreshold glutamatergic excitation (12 inputs). Gray trace: fast GABAergic synaptic activation (12 synapses each comprising 40 receptors with 25 pS peak conductance; Σg_GABA_ = 12 nS; PSP = 6.3 mV at dendritic spike site). Colored traces (blue to red): glutamatergic-GABAergic interaction at temporal offsets Δt = t_GLUT_ − t_GABA_ is illustrated from −30 to 150 ms; 10 ms intervals. (d) Summary of timing dependent effects on PSP amplitude. No effect of M_1_ receptor activation was observed. (e, f) As in (c, d) but for suprathreshold glutamatergic input. Fast synaptic GABAergic input reduced somatic output with only modest effects of M_1_ receptor mediated modulation. As in (c, d) but for suprathreshold glutamatergic input (15 inputs per site). Gray trace: fast synaptic GABAergic activation (40 synapses each comprising 40 receptors with 25 pS peak conductance; Σg_GABA_ = 40 nS). Colored traces (blue to red): glutamatergic-GABAergic interaction at temporal offsets Δt = t_GLUT_ − t_GABA_ of −100, −80, −60, −40, −20, respectively. Only a modest effect of M_1_ receptor activation on spike output was observed.

**Figure S5.**
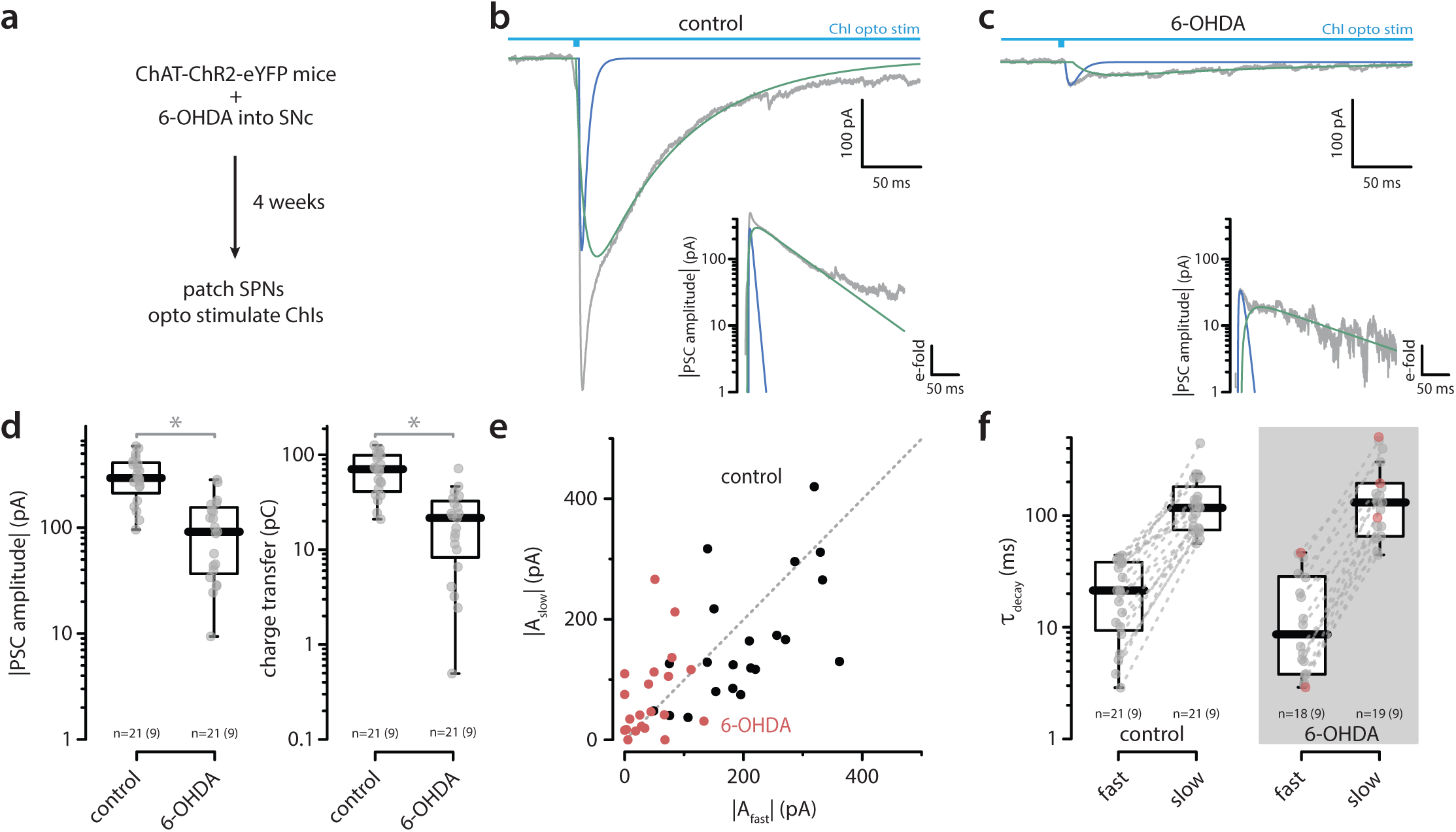
ChI-evoked GABAergic input in SPNs was reduced following 6-OHDA lesion in independent study. (a) Schematic of opsin injection into the DLS together with unilateral 6-OHDA injection into the substantia nigra pars compacta (SNc) of ChAT-ChR2-eYFP mice. Representative PSCs recorded from SPNs in control (b) and 6-OHDA (c) conditions in response to activation of ChIs (whole-field LED, 5 ms), showing fast- and slow-decaying gabazine-sensitive components. Insets, semi-logarithmic scale. (d) Semi-log box plots of PSC amplitude and charge transfer in SPNs (control: n = 21, 9 animals; 6-OHDA: n = 21, 9 animals) showing reduced responses after 6-OHDA (Wilcoxon rank-sum test, control vs 6-OHDA: amplitude, W = 39, p = 6.3 × 10^−7^; charge transfer, W = 404, p = 4.3 × 10^−7^). (e) Scatter plots of slow-decaying versus fast-decaying amplitudes in control (black) and 6-OHDA (red) conditions. (f) Semi-log box plots of fast and slow decay time constants in SPNs showing no change in kinetics (Wilcoxon rank-sum test, control vs 6-OHDA: fast, W = 240, *p* = 0.31292; slow, W = 188, *p* = 0.76828). As before, im some recordings, only one component (fast or slow) was identified (see Methods for the statistical criterion). These ‘unpaired’ decays are shown in red without a corresponding value in the other condition. Displayed n values are adjusted accordingly.

**Figure S6.**
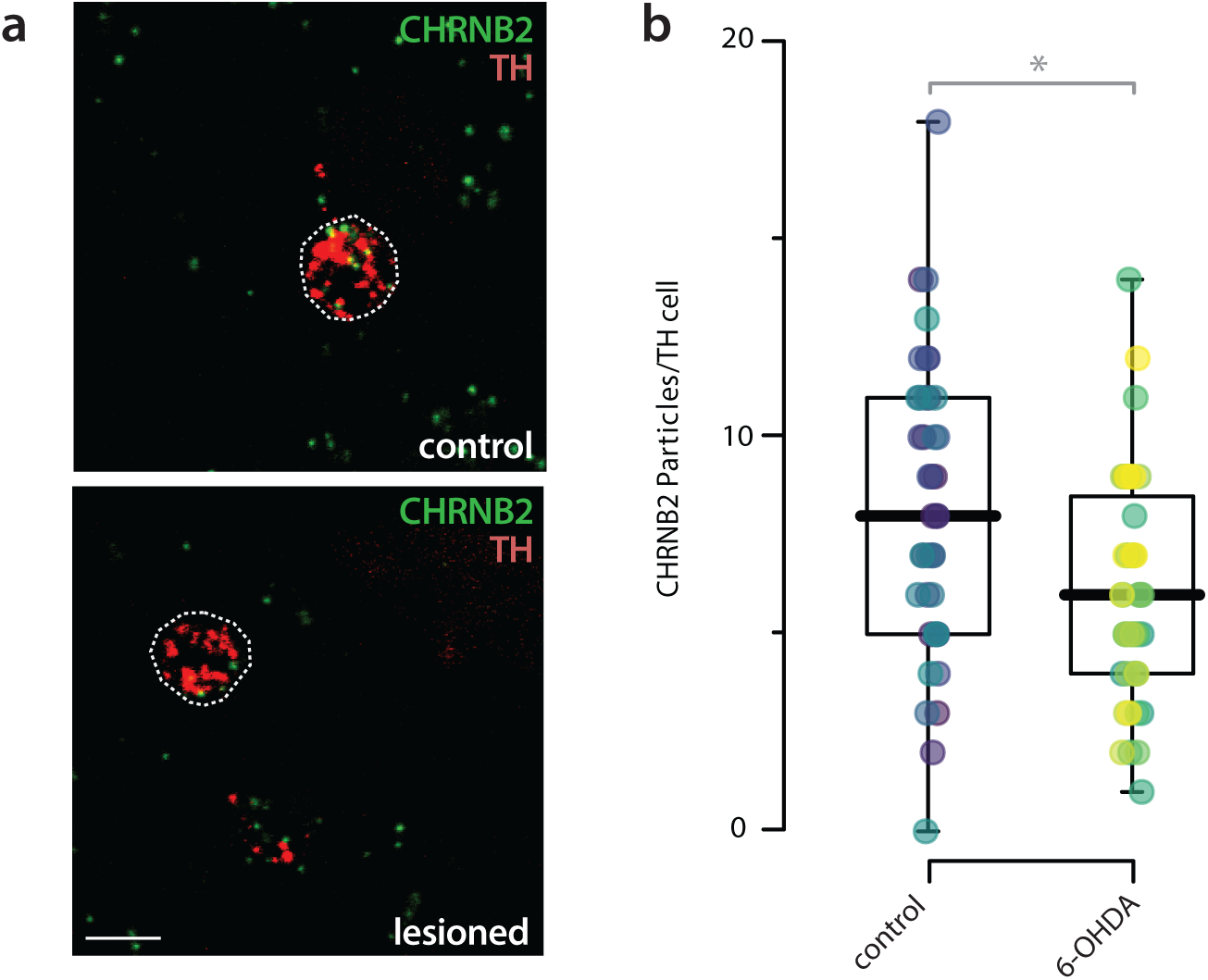
ChI-evoked input to TH+ neurons was reduced following 6-OHDA lesion. (a) In situ hybridization showing CHRNB2 (β2 subunit) mRNA expression in TH⁺ neurons under control and 6-OHDA conditions. (b) Box plots of β2-containing nicotinic acetylcholine receptor (nAChR) expression in TH⁺ neurons, demonstrating reduced expression following 6-OHDA (robust linear mixed-effects model, Nboot = 1,999, *p* < 0.0005).

